# The Presence of Periodontal Pathogens in Gastric Cancer

**DOI:** 10.1101/2020.03.23.003426

**Authors:** Marcel A. de Leeuw, Manuel X. Duval

**Affiliations:** GeneCreek, Inc.

## Abstract

**Background:** The microbiome is thought to play a role in the development of gastric cancer (GC). Several studies have put forward putatively carcinogenic species in addition to *Helicobacter pylori*, but are not in perfect alignment, possibly due to variable parameters in the experiments, including downstream processing. Meta-analyses have not been published so far, so there is lack in clinical guidance beyond *H. pylori* eradication therapy.

**Methods:** Here, we analysed gastric mucosa samples from nine public data sets, including GC samples. Using both unsupervised and supervised learning, we defined fine grain bacterial networks of gastric mucosa and identified species associated with tumor status of samples.

**Results:** We found anatomic locations and cohort regions among the possible factors leading to the observation of study specific gastric microbiomes. Despite this variability, the periodontal species *Fusobacterium nucleatum, Parvimonas micra* and *Peptostreptococcus stomatis* were found in association with tumor status in several datasets. The three species were predicted to be in interaction by ecological network analysis and also formed the intersection of tumor associated species between four GC data sets and five colorectal cancer (CRC) data sets we reanalyzed. We formulated a probiotic composition putatively competing with the GC pathogen spectrum, from correlation analysis in a large superset of gut samples (n=17,800) from clinical- and crowd-sourced studies.

**Implications:** The overlapping bacterial pathogen spectrum between two gastrointestinal tumor types, GC and CRC, has implications for etiology, treatment and prevention. In vitro testing results reported in literature suggest *H. pylori* eradication treatment should be efficient against the GC pathogen spectrum, yet the existence of an upstream periodontal reservoir is of concern. To address this, we propose longer term use of the formulated probiotics composition.

## Introduction

Gastric cancer (GC) is the sixth most common cancer in the world, with more than 70% of cases occurring in the developing world. GC is the third leading cause of cancer death worldwide (source: WHO, 2018). More than 50% of cases occur in Eastern Asia. In Asia, GC is the third most common cancer after breast and lung and is the second most common cause of cancer death after lung cancer [Rahman et al. 2014].

The seroprevalence of Helicobacter pylori is closely related to the incidence of GC [Kato et al. 2004, Ferreccio et al. 2007, Shiota et al. 2013]. In recent years, other bacteria have been proposed as risk factors for GC, including *Propionibacterium acnes* and *Prevotella copri* [Gunathilake et al. 2019], *Fusobacterium nucleatum* [Yamamura et al. 2017, Hsieh et al. 2018] and *Leptotrichia wadei* [Yang et al. 2016]. *Prevotella melaninogenica, Streptococcus anginosus* and *P. acnes* have been reported increased in the tumoral microhabitat [Liu et al. 2019]. The centrality of *Peptostreptococcus stomatis, S. anginosus, Parvimonas micra, Slackia exigua* and *Dialister pneumosintes* in GC tissue has been reported [Coker et al. 2018]. *P. acnes* has also been associated with lymphocytic gastritis [Montalban-Arques et al. 2016].

The availability of a number of these studies in the form of raw microbiome sequence reads offers the possibility to revisit the GC microbiome using a uniform bioinformatics approach, obtain a consensus of additional species possibly involved in GC and address ther-apeutic options beyond *H. pylori* eradication therapy.

## Materials & Methods

We identified a total of nine eligible datasets from literature and the NCBI BioProject repository. Exclusion criteria comprised the absence of quality data in the submission and the absence or mismatch of paired end sequences as submitted. Most eligible data sets are from China, Table 1. Scientific publication has been issued for the following projects: PRJEB21497 [Yap et al. 2016], PRJEB21104 [Parsons et al. 2017], PRJEB22107 [Klymiuk et al. 2017], PRJNA428883 [Liu et al. 2019] and PRJNA495436 [He et al. 2019]. For the purpose of comparison, we also included all five colorectal cancer (CRC) data sets we had previously analyzed, Table 2.

**Table 1:**
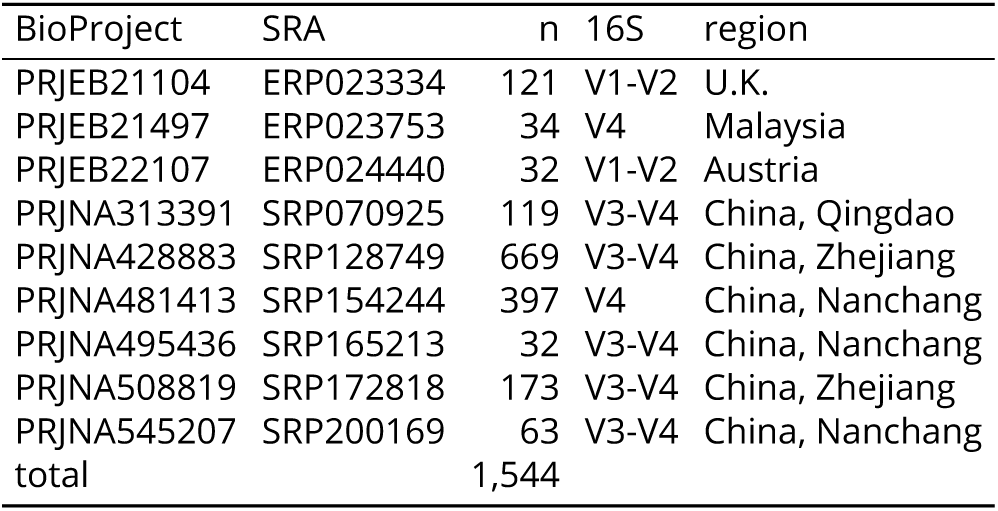
Gastric mucosa data sets used in this study. n: number of samples used, 16S: variable regions covered.

**Table 2:**
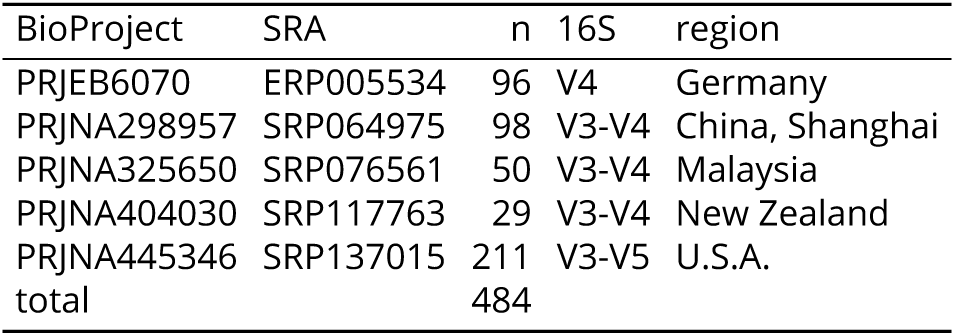
Colorectal cancer biopsy samples used in this study. n: number of samples used, 16S: variable regions covered.

| BioProject | SRA | n | 16S | region |
| --- | --- | --- | --- | --- |
| PRJEB6070 | ERP005534 | 96 | V4 | Germany |
| PRJNA298957 | SRP064975 | 98 | V3-V4 | China, Shanghai |
| PRJNA325650 | SRP076561 | 50 | V3-V4 | Malaysia |
| PRJNA404030 | SRP117763 | 29 | V3-V4 | New Zealand |
| PRJNA445346 | SRP137015 | 211 | V3-V5 | U.S.A. |
| total |  | 484 |  |  |

### Data analysis

Amplicon Sequence Variants (ASVs) were generated with the R Bioconductor package `dada2`, version 1.12.1 with recommended parameters [McMurdie, Paul J et al. 2016], involving quality trimming, discarding of sequences with N’s, assembly of forward and reverse sequences and contamination and chimera removal. ASVs per data set were subject to further analysis, involving multiple alignment with `mafft`, version 6.603b [Katoh et al. 2009] and approximatelymaximum-likelihood phylogenetic tree generation with `FastTreeMP`, version 2.1.11 [Price, Morgan N et al. 2010], both with default settings.

Taxonomic classification of ASVs were performed by `cur|sor` version 1.00, an in-house Python and R program using random forest (RF) based supervised learning on RDP release 11.5. The classifier assigns a species or higher level taxonomic identity to each ASV. Resulting classifications are available from the github repository https://github.com/GeneCreek/GC-manuscript in the form of R data objects.

UniFrac distances were computed using the R Bioconductor package `phyloseq`, version 1.28.0 [McMurdie and Holmes 2013] on raw ASVs. Further analysis used counts and relative abundances summarized at the species level, using the `cur|sor` provided taxonomic classifications.

Dirichlet Multinomial Mixtures (DMMs) were computed with the R bioconductor package `DirichletMultinomial`, version 1.26.0 [Holmes et al. 2012], using default parameters.

Downstream classification was performed using the R `caret` package, version 6.0.84, provided `rf` model. Variable (taxa) importance was estimated using the mean decrease in node impurity. Multiclass area-under-the-curve (AUC) [Hand and Till 2001] was computed by the R package `pROC`, version 1.15.3.

Ecological networks were computed using inverse covariance with SPIEC-EASI [Kurtz et al. 2015] as incorporated in the R Bioconductor package `SpiecEasi`, version 1.0.7, using default parameters.

For the nitrosating status of species, we required that at least one non-redundant genome for the species carries a UniProt annotated nitrate reductase alpha unit gene (narG) [Calmels et al. 1988].

Co-exclusion and co-occurrence between species for probiotics composition were computed using *χ*2 testing on detectable presence of species in samples (n=17,800) form a proprietary superset of over 40 clinical-and crowd sourced 16S studies, all performed on the Illumina platform.

## Results

### Pathogens in gastric mucosa

Among the species with highest prevalence in gastric mucosa of healthy individuals (n=85), we found a substantial number of opportunistic pathogens, with the majority being known as periodontal pathogens. Figure 1 depicts the distribution of prevalence and relative abundances of the top 20 periodontal and other pathogens. Whereas the position of *H. pylori* is obviously not a surprise, the 60% prevalence of the skin pathogen *P. acnes* (recently renamed to *Cutibacterium acnes*) is unexpected. The position of *F. nucleatum* among the top four pathogens is also remarkable. We found 17 distinct ASVs assigned to *P. acnes* and 53 distinct ASVs assigned to *F. nucleatum* in this dataset.

**Figure 1:**
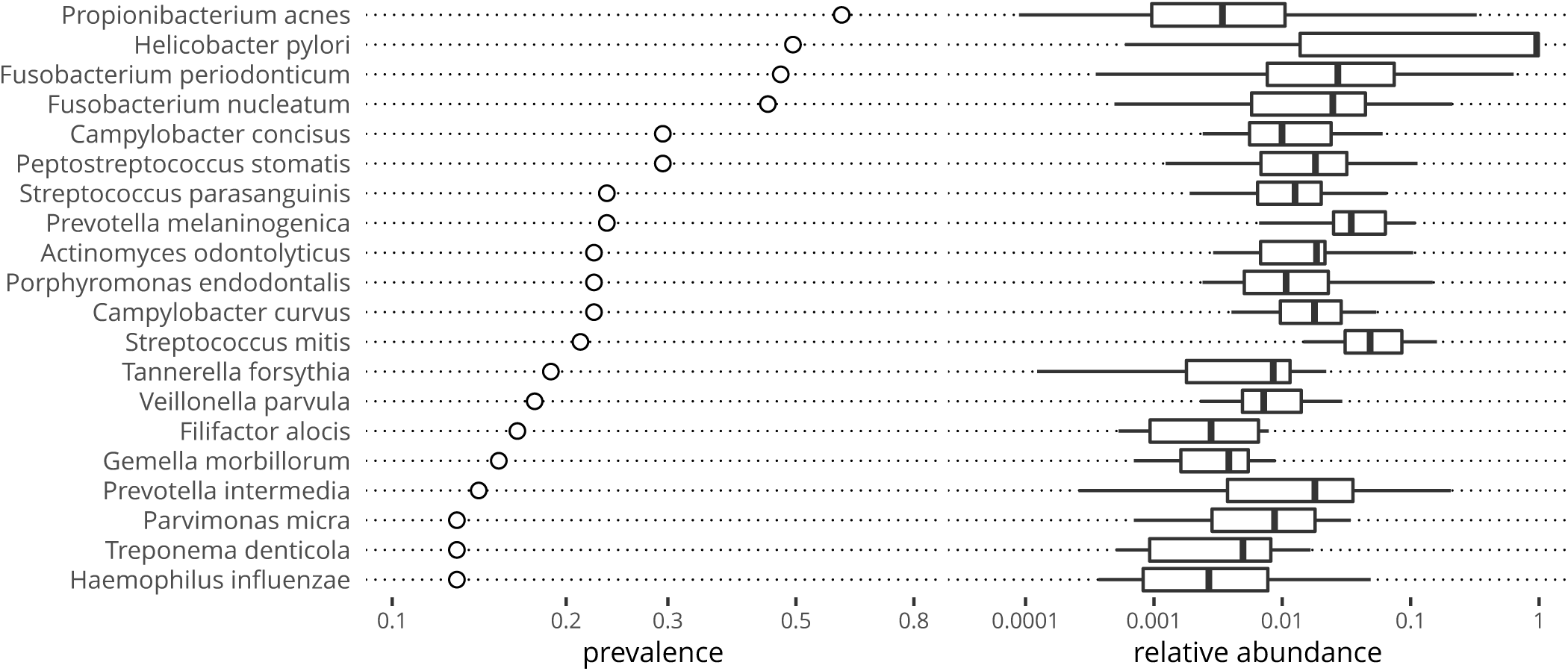
Distribution of prevalence and relative abundance of pathogens in healthy individuals.

### Gastric mucosa community analysis

We applied unsupervised clustering to investigate microbial gastric mucosa community structure, irrespective of sample disease status. In brief, using Dirichlet Multinomial Mixtures, we obtained an optimal goodness of fit at k=5 communities according to the Laplace and AIC evaluations, supplemental Fig. S1. Assigning per sample community types accordingly, we then retrieved the top 100 most important species. We assigned species to community types by maximum contribution. Putative interactions between these species were retrieved from the SPIEC-EASI ecological network constructor, which operated independently from the community structure on all 1,544 samples. Figure S2 depicts the correspondence between species community types and the correlation network.

For community types one and two the dominating species was *Helicobacter pylori*, with levels exceeding 50%, Fig. S3. Community type two had the lowest phylogenetic diversity of all community types, Fig. S4. Community type four received the majority of periodontal pathogens, whereas community types three and four harbored the most abundant nitrosating species, Table 3.

**Table 3:** Distribution of periodontal and other pathogens and nitrosating bacteria over community types.

| comm. type | periodontal | other | nitrosating |
| --- | --- | --- | --- |
| dmm 1 | 3 |  | 2 |
| dmm 2 |  | 1 |  |
| dmm 3 |  | 3 | 9 |
| dmm 4 | 20 | 5 | 8 |
| dmm 5 |  | 2 | 1 |

### Anatomical locations

Data set SRP154244 presents samples from different anatomical gastric locations in patients with gastritis, intestinal metaplasia and gastric cancer. We investigated if microbial signatures cluster by gastric location using random forest (RF) models and ecological networks, Table S3 and Fig. S5. Although we observed segregation between interacting antral curvature species on the one hand and corpus/antrum species on the other hand, it does not seem we can explain the distribution of data sets over the community types by difference in anatomical location alone.

### Disease progress

Data set SRP070925 contains gastric mucosa samples (n=119) from patients with gastritis, intestinal metaplasia, early gastric cancer and advanced gastric cancer. We combined this data set with data set SRP200169, containing gastric mucosa samples (n=63) from healthy subjects. Both are from Chinese cohorts and have been analysed using the 16S variable regions V3-V4 combined on the Illumina MiSeq. Performing multidimensional scaling on unweighted UniFrac distances, we found the disease stages are well separated, Fig. S6. We performed supervised learning of disease progress status with random forests on two thirds of the combined data set, with evaluation on the remaining third. Relative abundances summarized at the species level were used as the analysis substrate. Table 4 provides the classification results. Metaplasia were confounded with gastritis and early cancer, whereas advanced cancer samples were in part classified as early cancer. Healthy, gastritis and early cancer samples were well classified, resulting in an overall multi-class AUC of 0.936.

**Table 4:** Classification results on the disease stage evaluation subset, data set SRP070925. Predictions are in columns. Multiclass AUC:0.936.

| stage | healthy | gastritis | meta-plasia | early cancer | adv. cancer |
| --- | --- | --- | --- | --- | --- |
| healthy | 22 |  |  |  |  |
| gastritis |  | 10 |  |  |  |
| metaplasia |  | 4 | 2 | 3 |  |
| early cancer |  |  |  | 7 | 1 |
| advanced cancer |  |  |  | 5 | 7 |

### Sample disease location

Data set SRP128749 contains gastric mucosa samples (n=669) from GC patients and comprises triplet tumor, peripherical and normal samples. We added biopsies from healthy subjects to this cohort, again using data set SRP200169, to challenge the idea that GC normal reflects entirely healthy tissue. Performing multidimensional scaling on unweighted UniFrac distances, we found the disease locations show interesting separation, Fig. S9. We performed two supervised learning experiments on the combined data set, one with a two-thirds training, one-third evaluation setup and a second using one additional data set SRP172818 (n=173) also containing triplets as the cross-validation set. All three data sets are from Chinese cohorts and have been analysed using the 16S variable regions V3-V4 combined on the Illumina MiSeq.

Table 5 provides the classification results on the combined SRP128749 and SRP200169 data set. The model performs with a multi-class AUC of 0.842. Just one normal sample is confounded with healthy samples. The model performance increased to an AUC of 0.906 when trained on the whole combined data set and cross-validated on the SRP172818 data set, Table 6. None of the GC normal samples were confounded with samples from healthy donors.

**Table 5:**
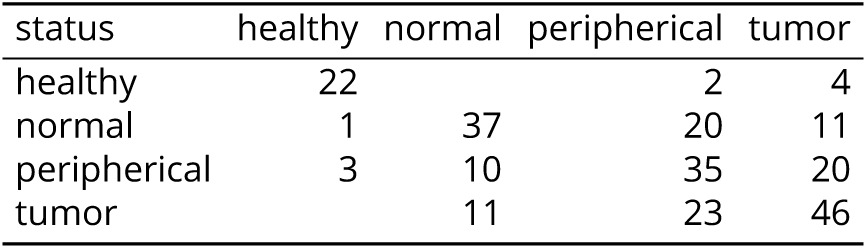
Combined SRP128749 and SRP200169 evaluation results. Predictions are in columns. Multiclass AUC:0.842

| status | healthy | normal | peripheral | tumor |
| --- | --- | --- | --- | --- |
| healthy | 22 |  | 2 | 4 |
| normal | 1 | 37 | 20 | 11 |
| peripheral | 3 | 10 | 35 | 20 |
| tumor |  | 11 | 23 | 46 |

**Table 6:** SRP172818 cross-validation results. Predictions are in columns. Multiclass AUC:0.906

| status | healthy | normal | peripheral | tumor |
| --- | --- | --- | --- | --- |
| healthy |  |  |  |  |
| normal |  | 45 | 8 | 4 |
| peripheral |  | 7 | 41 | 9 |
| tumor |  | 4 | 7 | 48 |

### Species relevant in GC

We disposed of four data sets having the metadata required for the association of species with tumor status, whether from a disease progress or disease location standpoint. In brief, we processed data sets individually and retrieved the top 50 differentiating species from the random forest models, trained on the data set as a whole. We generates ecological networks using these top species, retaining only connected nodes for display.

Figure 2 provides the putative interaction network of the disease location data sets SRP172818 and SRP128749, showing reproducible tumor association of, and possible interaction between, the oral species *F. nu-cleatum, P. micra, P. stomatis* and *Catonella morbi*. Correlation indicates the interaction would be cooperative.

**Figure 2:**
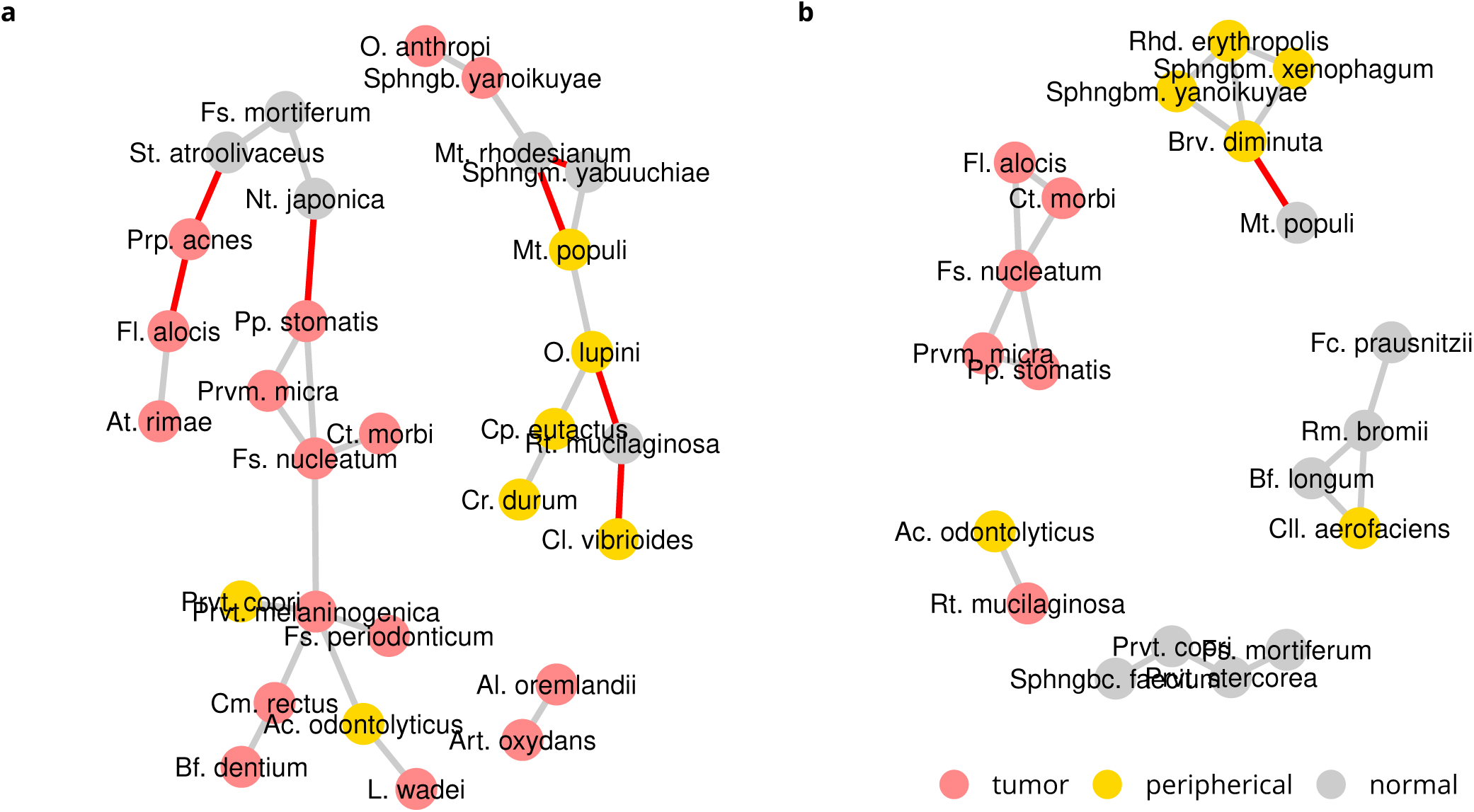
Disease status discriminating species. Data sets a) SRP172818 and b) SRP128749. Only species with interactions are displayed. Location associations are based on maximum mean relative abundance. Co-exclusion is indicated in red.

Supplemental Fig. S10 and Fig. S11 provide the same analysis for the disease progress data sets SRP070925 and ERP023334, respectively, in the first of which we found *P. melaninogenica* associated with advanced cancer status and in the second *F. nucleatum* with cancer status.

### Prevalence differences

An alternative take on the species differentiating between disease states, using *χ*2 testing of difference in prevalence, is presented in Tables S4-S8. *P. acnes* is reproducibly found at over 61% in GC tumor samples, whereas *P. stomatis* is found at over 54%, *P. micra* over 37% and *F. nucleatum* over 35% in GC tumor samples. The presence of all four roughly doubled over their baseline prevalence in normal samples, Tables S4 and S5.

### Comparison with colorectal cancer

We tested five previously analyzed colorectal cancer (CRC) data sets for presence and interactions of *F. nucleatum, P. micra* and *P. stomatis*. All five data sets SRP117763 (n=34, tumor-only) [Purcell et al. 2017], SRP137015 (n=211, tumor/peripherical/normal) [Hale et al. 2018b;a], SRP076561 (n=50, tumor/normal) [Drewes et al. 2017], ERP005534 (n=96, tumor/normal) [Zeller et al. 2014] and SRP064975 (n=98, tumor/peripherical/normal) [Lu et al. 2016] have been subject to publication. We found *F. nucleatum* in interaction with *P. stomatis* in SRP137015 and *P. micra* in interaction with *P. stomatis* in data sets SRP117763 and SRP076561, Fig. S12. Prevalence of *F. nucleatum* was found at 70% or more in tumor samples in SRP117763, Table S8, at 48% in tumor samples in SRP137015, Table S9 and at 73% in tumor samples in SRP076561, Table S10. Listing the most abundant cancer associated species in GC and CRC, the intersection between the two cancer types was formed by *F. nucleatum, P. micra* and *P. stomatis*, Table 7.

**Table 7:** Correspondence between GC- and CRC-associated species. Numbers reflect the number of datasets in which the species is found associated, out of four possible. Species found in more than one dataset and with relative abundance > 0.5% in cancer are listed.

| species | GC | CRC |
| --- | --- | --- |
| *Bacteroides fragilis* |  | 2 |
| *Bacteroides ovatus* |  | 3 |
| *Brevundimonas vesicularis* | 2 |  |
| *Escherichia coli* |  | 2 |
| *Fusobacterium nucleatum* | 3 | 3 |
| *Gemella morbillorum* |  | 3 |
| *Parvimonas micra* | 2 | 3 |
| *Peptostreptococcus stomatis* | 2 | 2 |
| *Prevotella intermedia* |  | 2 |
| *Propionibacterium acnes* | 2 |  |

QPS species

### Eradication therapy

Data set SRP165213 provides mucosa samples, pre- and post bismuth quadruple *H. pylori* eradication therapy. Using *χ*2 testing of difference in prevalence, we found several bacteria, including the expected *H. pylori*, exhibit an important drop in prevalence, Table 8. *P. stomatis, P. micra* and *F. nucleatum* on the other hand showed a moderately significant prevalence increase.

**Table 8:**
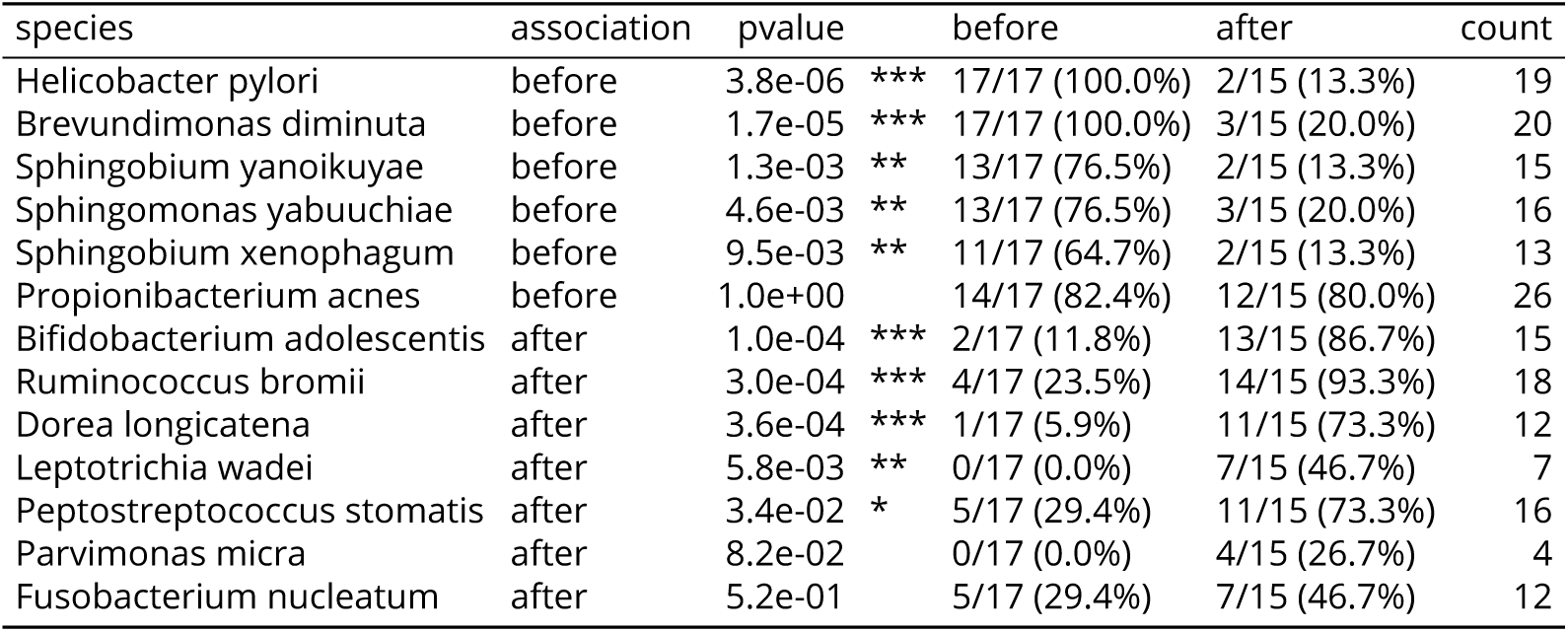
prevalence differences between before and after Hp eradication, SRP165213.

### Modulation of the gastric mucosa microbiome

Using prevalence data from 17,800 gut samples, including the samples used in this study, we probed for qualified presumption of safety (QPS) species found in co-exclusion with the species of interest panel identified above. Figure 3 shows the result. *Bifidobacterium longum* appears as the most promising QPS species, followed by *Streptococcus salivarius* both of which are being used in probiotic products and are actually detectable in gastric mucosa samples, see Fig. 3b for *B. longum*. In the healthy data set SRP200169 we found 27 ASVs for *B. longum* but none for *S. salivarius*, indicating that the latter is probably not commensal in the stomach in healthy individuals.

**Figure 3:**
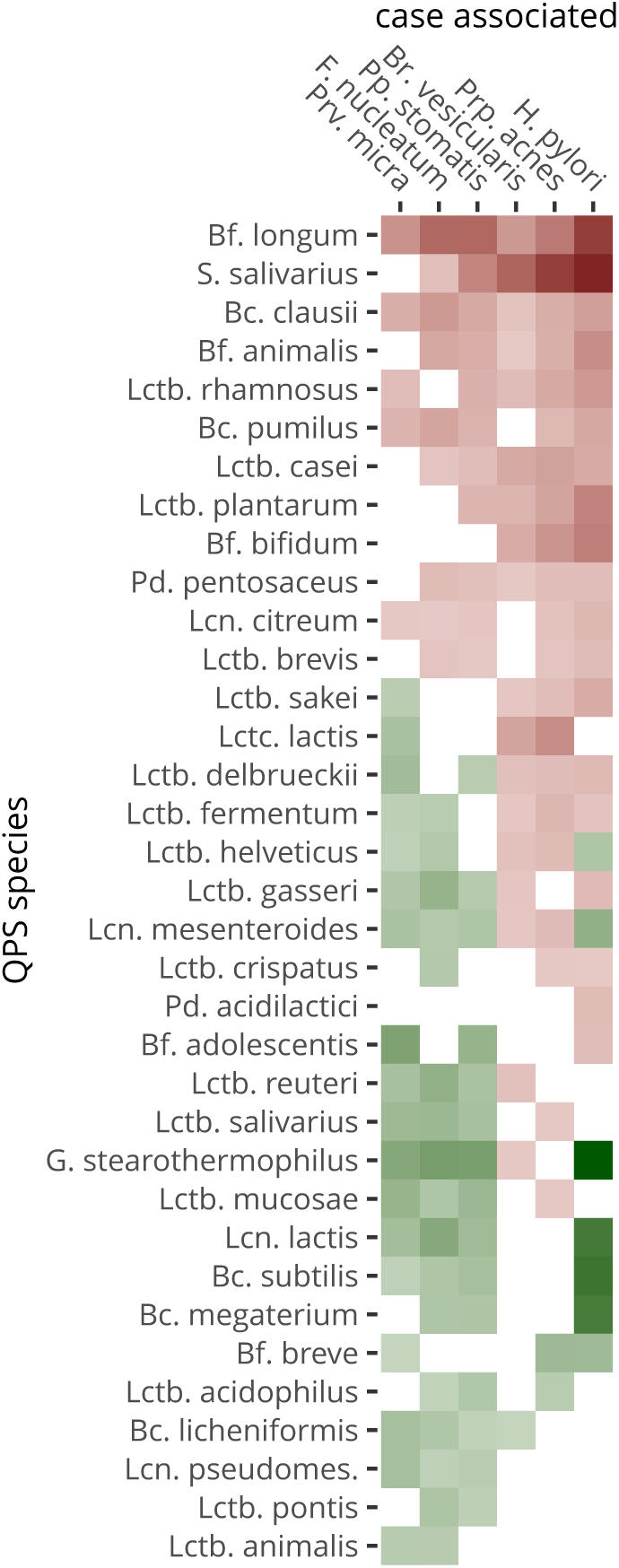
Co-exclusion by and co-occurrence with QPS species of GC associated species. Putative inhibition is in shades of red, potential synergy in shades of green. White reflect neutrality or too little combined prevalence to make a call. Genera are abbreviated as follows: **Bcl.**: *Bacillus*, **Bf.**: *Bifidobacterium*, **Gb.**: *Geobacillus*, **Lcn.**: *Leuconostic*, **Lctb.**: *Lactobacillus*, **Lctc.**: *Lactococcus*, **Pd.**: *Pediococcus*, **S.**: *Streptococcus*.

## Discussion

In this above, we revisited public gastric mucosa and colorectal cancer data sets, taking into account recent advances in processing of amplicon metagenomic sequences [Callahan et al. 2017], establishing species level taxonomic classification.

### Limitations

Use of a healthy cohort analyzed as a separate batch and from a different regional cohort does not allow to control for batch- or regional effects in supervised learning. Regional clustering of GC microbiota has been reported previously [Yu et al. 2017]. So our case that samples from healthy donors are distinct from GC normal samples in GC patients is a delicate case. For confirmation of this hypothesis, healthy donors need to be recruited from the same population as the GC patients.

Four subspecies are known for *F. nucleatum*. Our taxonomic classifier does not resolve down to the level of subspecies, so all counts and relative abundances for *F. nucleatum* may conceal different subspecies, moreover so since in CRC, multiple subspecies have been isolated from biopsies [Brennan and Garrett 2019] and since we detected several tens of distinct ASVs associated with *F. nucleatum*.

### Low biomass and contamination

*P. acnes* has been proposed as a possible contaminant of many experiments [Mollerup et al. 2016]. This is particularly relevant for gastric samples which are of low biomass as compared to biopsies from the lower GI tract. That does not mean we need to discard the bacterium altogether, notably not if it shows significant increase in tumor sample locations as in data sets SRP172818 and SRP128749, but it could mean its baseline presence is overestimated and hence its status as a gastric mucosa commensal [Delgado et al. 2011]. Its position as a prevalent but low abundant species in healthy subjects gives credit to the contamination thesis. The number of ASVs associated with *P. acnes* though suggests that if there is contamination, it originates from multiple individuals. The fact that the bacterium never reached high abundance in the experiments means that it did not contaminate low biomass samples in particular.

### Helicobacter pylori

In all data sets, we found gastric mucosa samples completely exempt of H. pylori, including in normal and peripherical samples, which opens the possibility that other pathogens play a role in GC. We did not find *H. pylori* in significant interaction, which is unexpected and discrepant to findings on the same data set SRP128749 reported [Liu et al. 2019]. We attribute this discrepancy to the use of a more stringent ecological network inference [Kurtz et al. 2015]. On the other hand, report has been made that *H. pylori* presence did not affect microbial community composition [Bik et al. 2006]. So it seems that although *H. pylori* may create oncogenic conditions through host interaction, there does not seem to be a direct benefit or detriment of such conditions for other bacteria.

### Cohort specific species

Our results show species found in gastric mucosa have a strong cohort specific distribution of species. Within cohort prediction of sample disease status or location status based on the microbiome composition is performing well with AUCs over 0.8, so despite its diversity, there is a clear sample status signature in the microbiome composition.

### Nitrosating species

Nitrosating bacteria convert nitrogen compounds in gastric fluid to potentially carcinogenic N-nitroso compounds (NOCs), which are believed to contribute to gastric cancer [Sharma et al. 1984, Mowat et al. 2000, Jo et al. 2016, Ferreira et al. 2018, Park et al. 2019]. We found nitrosating bacteria were not uniformly distributed over gastric mucosa community types. Community type four combines nitrosating species with periodontal pathogens and can be considered as the highest GC risk community type.

### Periodontal and CRC pathogens

It has been reported that among patients with periodontal disease, high levels of colonization of periodontal pathogens are associated with an increased risk of gastric precancerous lesions [Salazar et al. 2013]. We found the periodontal pathogens *F. nucleatum, P. micra* and *P. stomatis* to be commensal but also associated with tumor status and in direct interaction in several data sets. These three species were also found in association with tumor status in CRC data sets revisited and correspond with a CRC subtype with strong immune signature [Purcell et al. 2017]. Revisiting the CRC data sets, we found in part the same interactions as in GC. Two recent meta-analysis of CRC case-control studies placed *F. nucleatum, P. micra* and *P. stomatis* among the top five carcinoma enriched species [Drewes et al. 2017, Wirbel et al. 2019]. *F. nucleatum* and *P. stomatis* have also been proposed among a panel of species for early detection of CRC [Zeller et al. 2014].

### Virulence

The gram negative bacterium *F. nucleatum* promotes tumor development by inducing inflammation and host immune response in the CRC microenvironment. Its adhesion to the intestinal epithelium can cause the host to produce inflammatory factors and recruit inflammatory cells, creating an environment which favors tumor growth. Treatment of mice bearing a colon cancer xenograft with the antibiotic metronidazole reduced Fusobacterium load, cancer cell proliferation, and overall tumor growth [Bullman et al. 2017]. *F. nucleatum* can induce immune suppression in gut mucosa, contributing to the progression of CRC [Wu et al. 2019]. In CRC, *F. nucleatum* is predicted to produce hydrogen sulfide (*H*_2_*S*) [Hale et al. 2018b], which is a metabolite with a dual role, both carcinogenic and antiinflammatory. Epithelial cells react to *F. nucleatum* by activation of multiple cell signaling pathways that lead to production of collagenase 3, increased cell migration, formation of lysosome-related structures, and cell survival [Uitto et al. 2005].

Furthermore, it is predicted *F. nucleatum* infection regulates multiple signaling cascades which could lead to up-regulation of proinflammatory responses, oncogenes, modulation of host immune defense mechanism and suppression of DNA repair system [Kumar et al. 2016]. There does not seem to be a reason why *F. nucleatum* would not be pathogenic in gastric tissue whereas it is in periodontal, respiratory tract, tonsils, appendix, colonic and other tissues [Han 2015].

The gram positive anaerobe *P. stomatis* has been isolated from a variety of periodontal and endodontic infections, as well as infections in other bodyparts [Downes and Wade 2006]. The species has been found associated with oral squamous cell carcinoma (OSCC) [Pushalkar et al. 2012]. At present, little is known about the specifics of its pathogenicity. The type strain (DSM 17678) genome harbors a gene (mprF, phosphatidylglycerol lysyltransferase) producing lysylphosphatidylglycerol (LPG), a major component of the bacterial membrane with a positive net charge. LPG synthesis contributes to bacterial virulence as it is involved in the resistance mechanism against cationic antimicrobial peptides produced by the host’s immune system and by competing microorganisms. Contrary to other *Peptostreptococci, P. stomatis* does not produce intestinal barrier enforcing indole-3-propionic acid (IPA) or indoleacrylic acid (IA) [Wlodarska et al. 2017].

*P. micra*, previously known as *(Pepto)streptococcus micros*, is a gram positive anaerobe which is known to be involved in periodontal infections. It has also been isolated from OSCC [Hooper et al. 2007]. It is a producer of collagenase and of limited elastolytic and hemolytic activity [Ota-Tsuzuki and Alves Mayer 2010]. In a mouse CRC model, *P. micra* elicited increased Th2 and Th17 cells, decreased Th1 cells and increased inflammation [Yu et al. 2019].

### The oral cavity as reservoir

It has been shown that in a number of cases (6/14, 43%) identical *F. nucleatum* strains could be recovered from CRC and saliva of the same patients [Komiya et al. 2019]. Furthermore, the oral microbiome composition is to a certain extent predictive for CRC disease progress status [Flemer et al. 2018]. It is tempting to speculate that a similar relationship could be explored for disease progress in GC.

### Biofilm formation

*F. nucleatum* is regarded as a central organism for dental biofilm maturation due to its wide ability to aggregate with other microorganisms, such as *Porphyromonas gingivalis* [Tavares et al. 2018]. It is considered as a bridge bacterium between early and late colonizers in dental plaque [He et al. 2016]. The eventuality of *H. pylori*- and non *H. pylory* biofilm formation in the gastric environment has been raised [Rizzato et al. 2019]. Our ecologic interaction networks suggests *F. nucleatum* and other bacteria, but not *H. pylori*, could indeed engage in gastric mucosa biofilms and more particularly in GC biofilms.

### Antibiotherapy

*Helicobacter pylori* eradication therapy has been shown to have a prophylactic effect against GC [Kwok et al. 2008]. The first-line therapy consists of a proton pump inhibitor (PPI) or ranitidine bismuth citrate, with any two antibiotics among amoxicillin, clarithromycin and metronidazole. In vitro testing has shown *Peptostreptococcus stomatis* is sensitive to amoxicillin and metronidazole [Könönen et al. 2007]. *F. nucleatum* is sensitive to amoxicillin or amoxicillin/clavulanate combination therapy [Jacinto et al. 2008] and to metronidazole [Shilnikova and Dmitrieva 2015, Bullman et al. 2017]. *Parvimonas micra* is sensitive to amoxicillin/clavulanate and metronidazole [Veloo et al. 2011]. In vivo sensitivity of the species may differ and in addition, with the oral cavity as a reservoir, periodontal pathogens could recolonize the gastric environment and take advantage of the space cleared by *H. pylori*, which is what our data suggests.

### Probiotics use

We predicted in silico that several QPS species could be effective against the spectrum of *H. pylori* and the periodontal pathogens discussed above. Our findings are coherent with the report that probiotics including *Bifidobacterium longum, Lactobacillus acidophilus*, and *Enterococcus faecalis* significantly reduced the abundance of F. nucleatum in CRC surgery patients by nearly 5-fold, whilst normalizing dysbiosis [Gao et al. 2015]. In vitro adhesion inhibition of gram-negative species by *B. longum* has been reported [Inturri et al. 2016]. Other than adhesion inhibitors, bifidobacteria produce acetate and lactate as well as vitamins, antioxidants, polyphenols, and conjugated linoleic acids which have been proposed to act as chemical barrier against pathogen proliferation [Inturri et al. 2019]. *Streptococcus salivarius* not only inhibits adhesion of pathogens to epithelial cells, but also produces bacteriocins [Manning et al. 2016].

## Prospects

In future GC microbiome studies, it appears imperative to include normal controls from healthy subjects so that normal samples from GC patients can be properly compared against samples from healthy subjects. Fluorescent in situ hybridization could be used in case of gastrectomy to confirm biofilm status of the aforementioned pathogen spectrum. A long term maintenance formula using probiotics after an antibiotics eradication course can be of interest as a treatment option. A variety of *Bifidobacterium longum* strains are used in several probiotic preparations commercially available whereas *Streptococcis salivarius* strain K12 [Burton et al. 2006] is also commercially available.

## Conclusions

In conclusion, we found disease progress and sample disease status is not reflected in the overall bacterial community type of mucosa. Rather, community types are populated by potentially regionally distinct species. Despite this diversity, we found periodontal pathogens as a common denomicator. These pathogens were also identified in CRC, establishing possible microbial similarities between subtypes of GC and CRC, with implications for etiology, treatment and prevention. Correlation networks suggest these species, as in dental plaque and in CRC, engage in biofilm formation in gastric mucosa. Probiotics should be considered as a treatment option, after *H. pylori* eradication therapy, to avoid recolonization by periodontal pathogens.

## Acknowledgements

The authors received no financial support for this study. The authors acknowledge the contributions to the Short Read Archive made by the respective institutions and acknowledge scientific journals for enforcing this practice.

## Declarations

### Authors’ contributions

Study design, data collection, data analysis and writing of the manuscript (ML); data analysis and writing of the manuscript (MD).

