## supplemental materials for "The Presence of Periodontal Pathogens in Gastric Cancer"

### Supplemental analyses

#### microbial community types

Using Dirichlet Multinomial Mixtures on the combined relative abundances of the nine datasets (n=1,544) listed in table 1, we obtain an optimal goodness of fit at k=5 communities according to the Laplace and AIC evaluations, figure S1. The breakdown of samples from the various datasets along the community types is given in table S1.

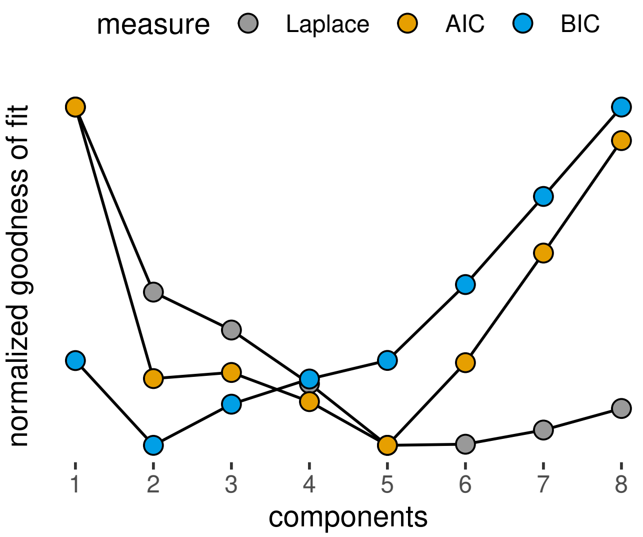

**Figure S1:** Goodness of fit of the DMM models at different k.

**Table S1:** Distribution of community types across studies. The five community types are in columns.

| study | dmm 1 | dmm 2 | dmm 3 | dmm 4 | dmm 5 |
|---|---|---|---|---|---|
| ERP023334 | 10 | 30 | | 81 | |
| ERP023753 | 16 | 5 | | 13 | |
| ERP024440 | 1 | 13 | | 18 | |
| SRP070925 | 2 | | | | 117 |
| SRP128749 | 635 | 34 | | | |
| SRP154244 | | 83 | 179 | 39 | |
| SRP165213 | 23 | 9 | | | |
| SRP172818 | 155 | 17 | | 1 | |
| SRP200169 | 42 | 21 | | | |

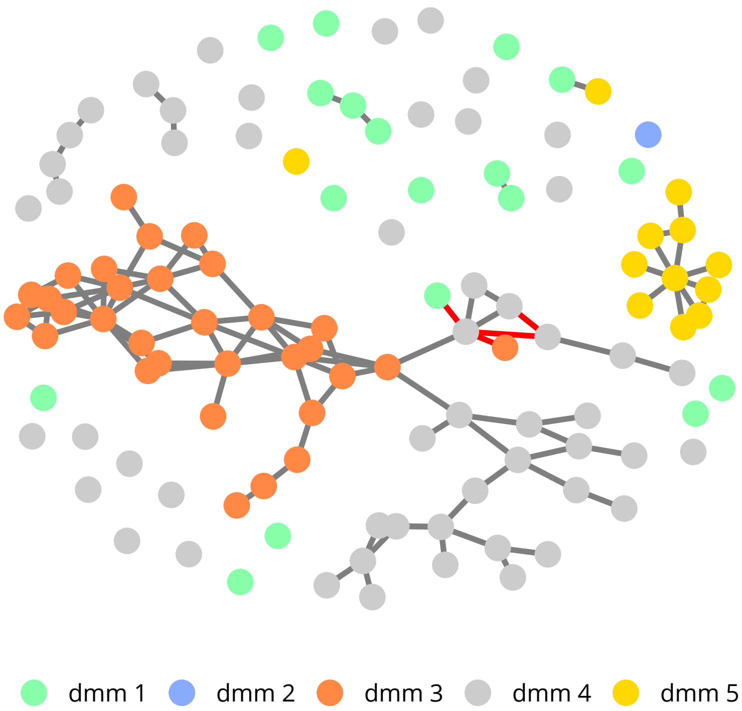

**Figure S2:** Interaction network between species relevant for community types. The top 100 species relevant for distinction between the five community types are displayed.

The first two community types are dominated by a few species mostly without interaction. The majority of healthy donor samples was located in community type one, together with certain tumor samples.

Of note, community types three and five received contributions from a single study each, Table S1. Hence, although we find multinomial mixtures and inverse covariance networks were in good agreement for overall gastric microbiota composition, we observed potentially only a subset of regionally or otherwise determined gastric microbiota. Among the top 100 differentiating species we found 62 distinct genera, further highlighting the diversity. Table S2 lists the 18 genera with more than one species.

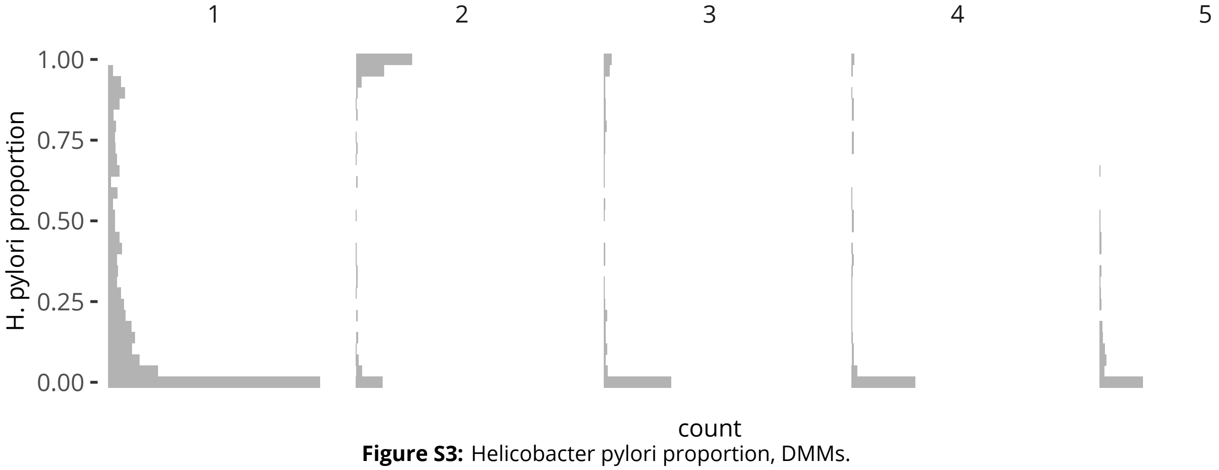

**Figure S3:** Helicobacter pylori proportion, DMMs.

**Table S2:** Gastric mucosa genera. Only genera with more than one species are listed.

| genus | species |
|---|---|
| Prevotella | 10 |
| Streptococcus | 9 |
| Acinetobacter | 4 |
| Campylobacter | 4 |
| Porphyromonas | 4 |
| Arthrobacter | 3 |
| Fusobacterium | 3 |
| Leuconostoc | 3 |
| Methylobacterium | 3 |
| Sphingomonas | 3 |
| Veillonella | 3 |
| Actinomyces | 2 |
| Alloprevotella | 2 |
| Bacillus | 2 |
| Brevundimonas | 2 |
| Clostridium | 2 |
| Haemophilus | 2 |
| Lactococcus | 2 |
| Neisseria | 2 |

Further indication that the DMMs are distinct in nature can be found in the projection of alpha diversity, using the phylogenetic diversity (whole tree), figure S4. The *Helicobacter pylori* dominated community type three has the lowest diversity.

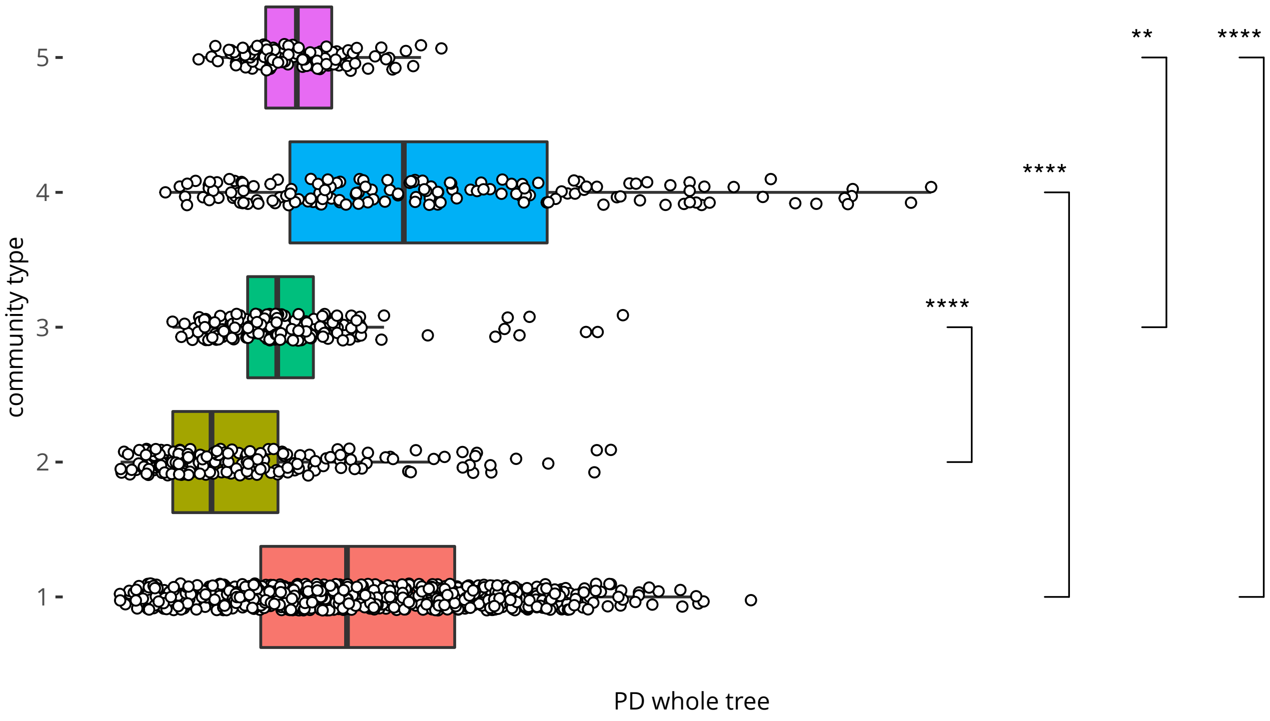

**Figure S4:** Alpha diversity of community types. Phylogenetic diversity (whole tree).

#### anatomic location

Data set SRP154244 presents samples from different gastric locations in patients with gastritis, intestinal metaplasia and gastric cancer. We investigate if microbial signatures differ per anatomic location by training an RF model on two thirds of the samples and evaluating the model on the remaining third. Table S3 suggests the antral is well differentiated from the antrum and body, but the latter two are not differentiated. Thus at first sight, gastric location could at least in part explain differences in community types.

**Table S3:** RF classification of sampling location. Predictions are in columns. Multiclass AUC:0.788

| location | antral | antrum | body |
|---|---|---|---|
| antral | 72 | 3 | 0 |
| antrum | 6 | 11 | 0 |
| body | 1 | 6 | 1 |

To shine further light on this matter, we group corpus and antrum samples together and retrain an RF model on the whole of the SRP154244 dataset, retrieve differentiating species and build a SPIEC-EASI network, figure S3. Although we find significant separation between the two locations, especially when considering the negative correlations (in red), the separation is not as strict as the separation between community types. So it does not seem we can explain the distribution of datasets over the community typrs by difference in anatomic location alone. Of note, we find three bacteria encountered in colorectal cancer, *Fusobacterium nucleatum*, *Parvimonas micra*, *Peptostreptococcus stomatis* in interaction and associated with the corpus/antrum. *Helicobacter pylori* is more abundant in the antral and not in interaction with any other species.

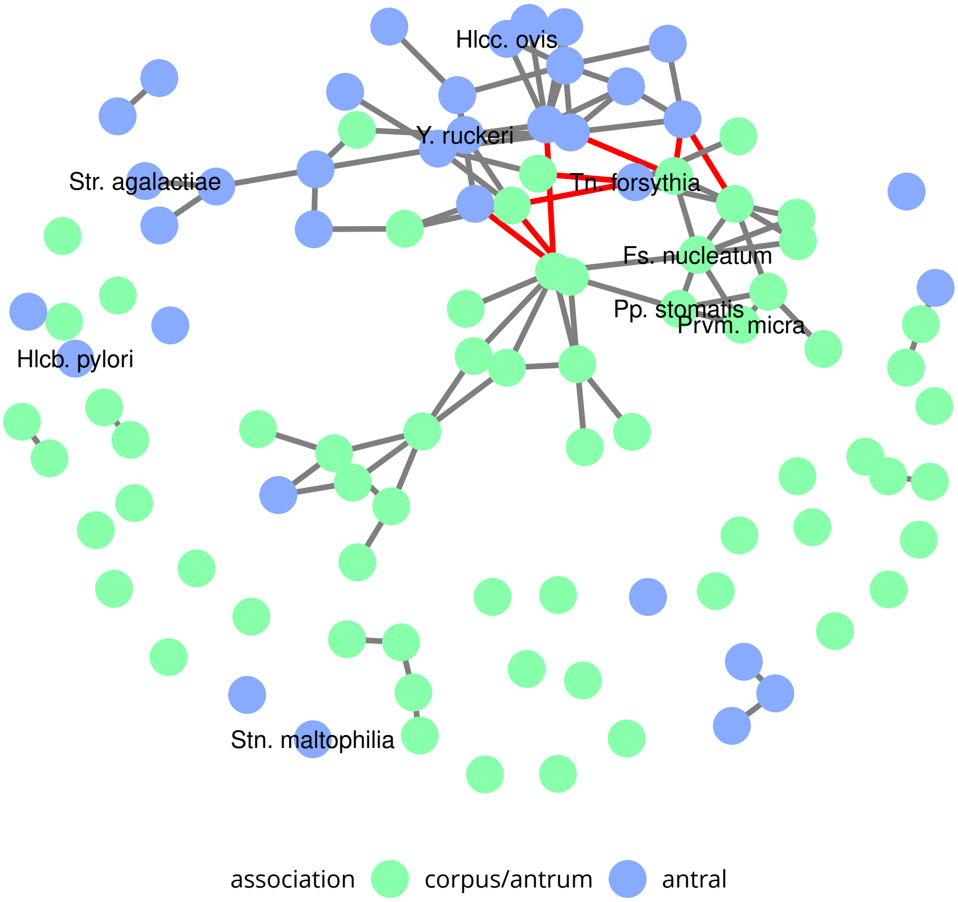

**Figure S5:** Interaction network between species relevant for gastric location. The top 100 species relevant for distinction between the two gastric locations are displayed. Opportunistic pathogens are labelled.

#### disease progress

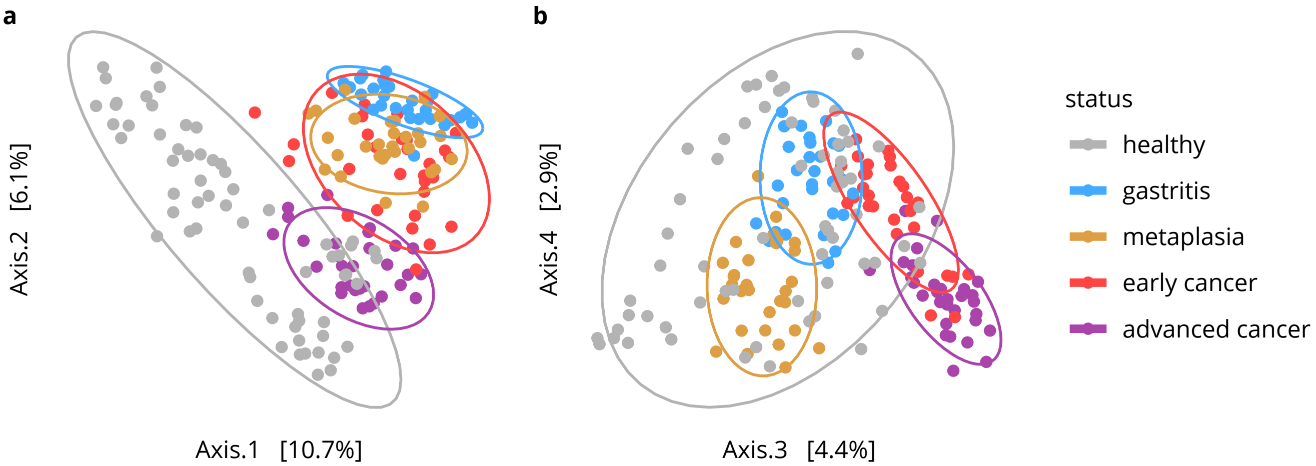

**Figure S6:** Multi-dimensional scaling of the disease progress data set. Unweighted UniFrac of ASVs is used as the distance metric.

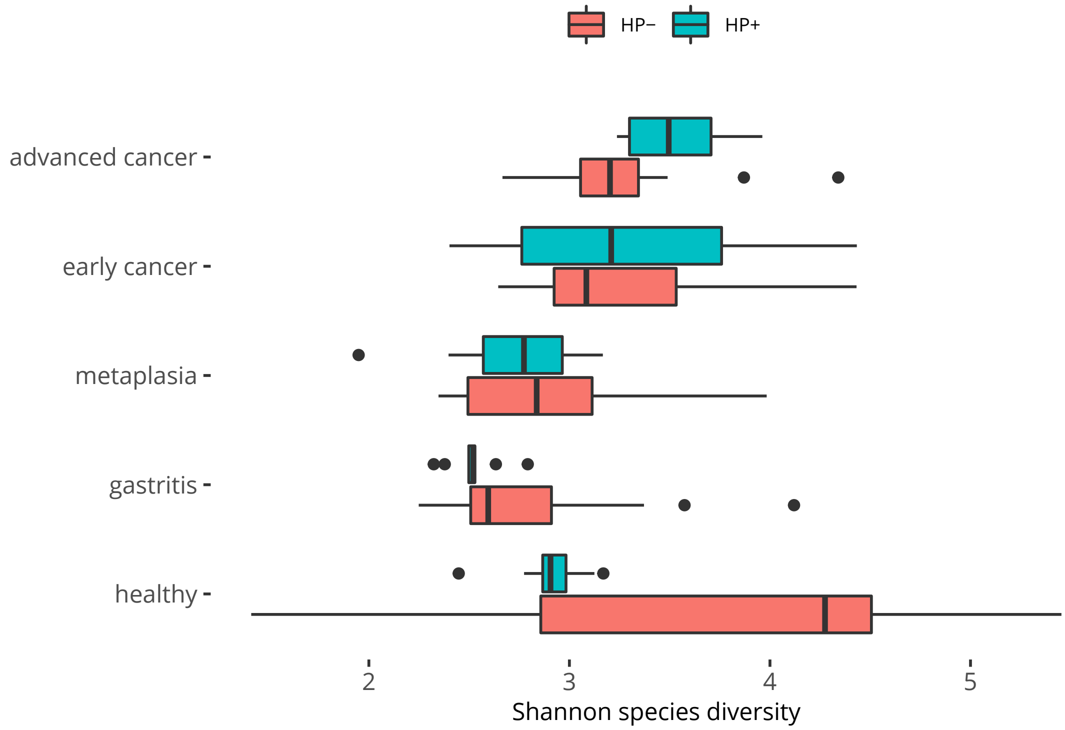

**Figure S7:** Shannon species diversity and disease progress (SRP070925). Helicobacter pylori positive (Hp+) and negative (Hp-) samples are distinguished.

The alpha diversity evolves along the disease progress path according to the Shannon criterium, figure S7 and S8. The gastritis is characterized by dysbiosis as compared to healthy tissue, with a trend to reach normal diversity along the disease progress.

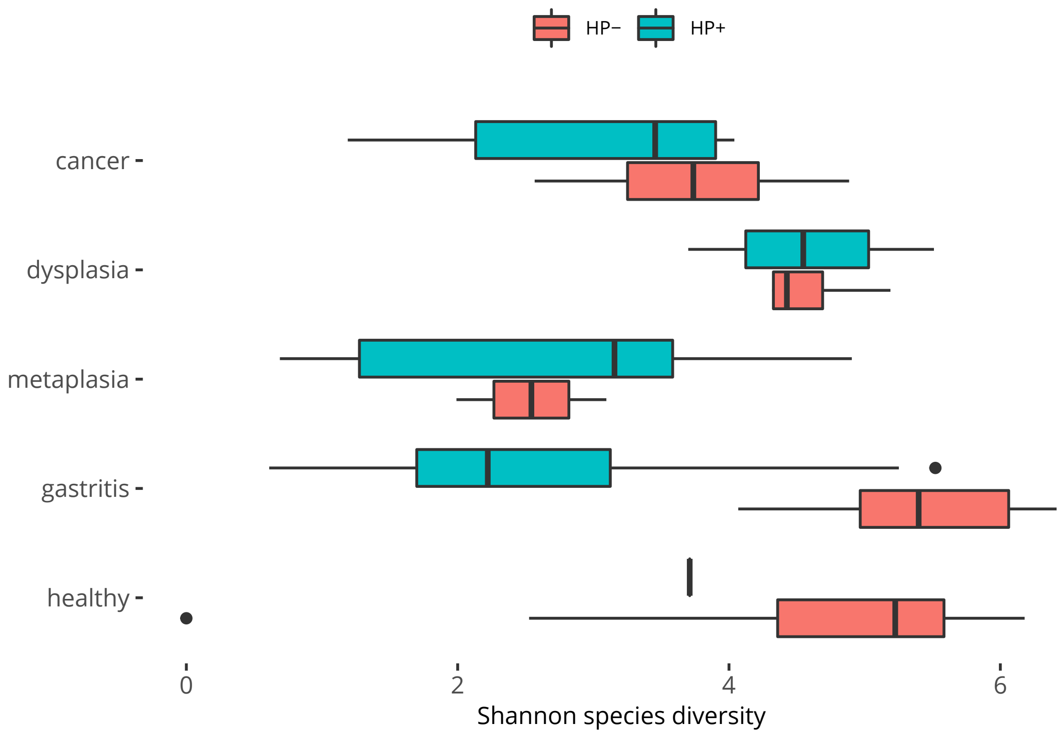

**Figure S8:** Shannon species diversity and disease progress (ERP023334). Helicobacter pylori positive (Hp+) and negative (Hp-) samples are distinguished.

##### disease location

Using unweighted UniFrac distance on ASVs (amplicon sequence variants) we obtain better MDS separation of normal/peripherical/tumor samples than reported in [Liu et al. 2019], using the same dataset, whether without (not shown) or with addition of samples from healthy donors, figure S4.

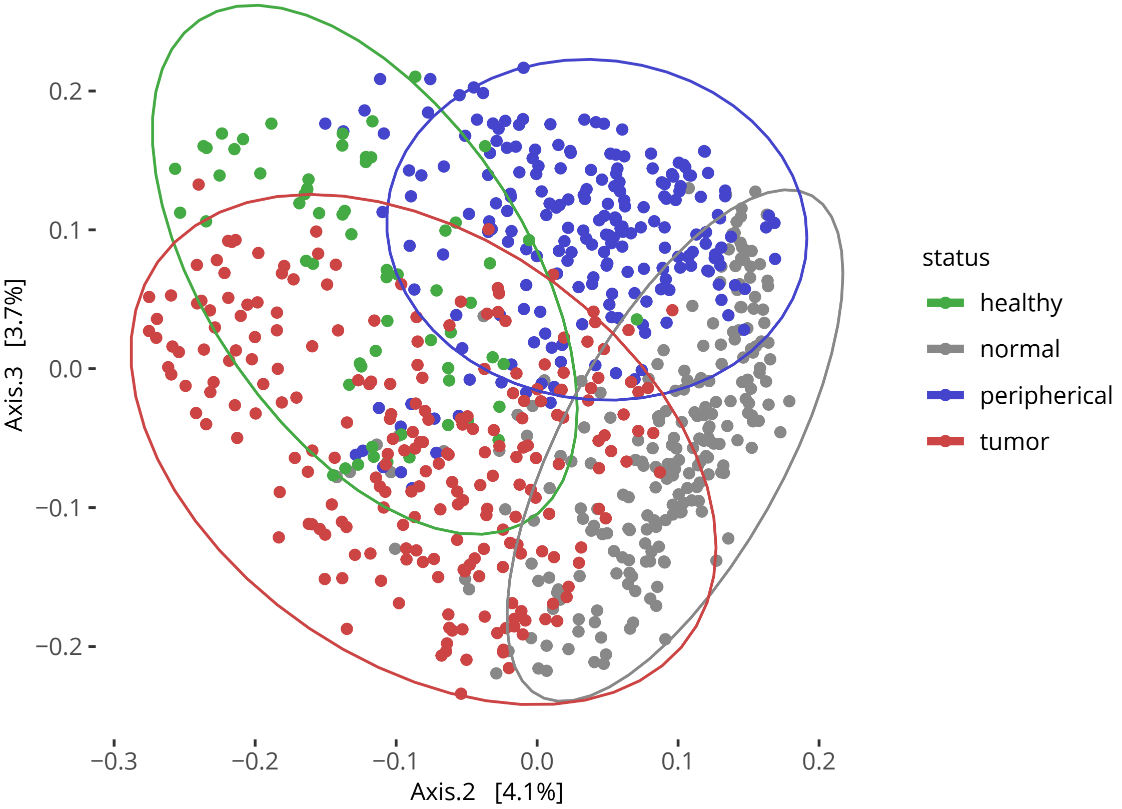

**Figure S9:** Multi-dimensional scaling of the disease status dataset SRP128749. Unweighted UniFrac of ASVs is used as the distance metric.

**relevant species in GC**

We dispose of four datasets with the metadata required for the association of species with tumor status, whether from a disease progress or tumor/normal status standpoint. We choose to process datasets individually because of possible regional differences and retrieve the top 50 differentiating species from the random forest models, which we train on the datasets as a whole, so as to maximize performance. We provide sequence counts of these top 50 species to Spiec Easi for ecological network generation. We retain only connected nodes for display. Figure 3 provides the result for the two tumor/peripherical/normal datasets SRP128749 and SRP172818 alongside for comparison. Figure S10 below provides the same for the disease progress data set SRP070925, figure S11 for the disease progress data set ERP023334.

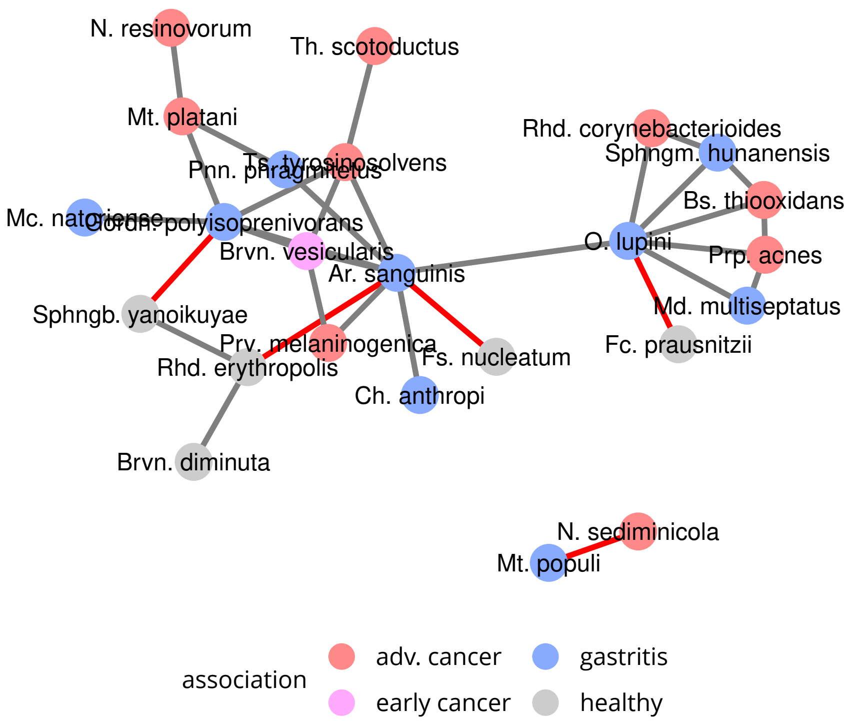

**Figure S10:** Interaction network between species relevant for disease progress (SRP070925). The top 50 species relevant for distinction between healthy the three disease stages are displayed. Only species with interactions are shown. Co-exclusion interactions are displayed in red.

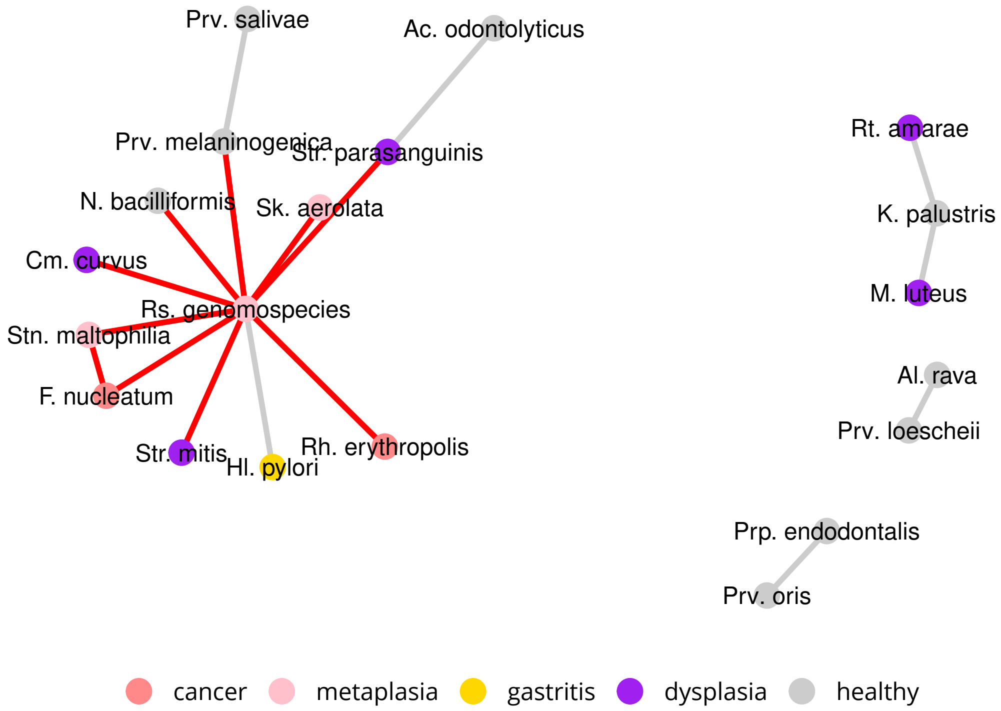

**Figure S11:** Interaction network between species relevant for disease progress (ERP023334). The top 50 species relevant for distinction between healthy four disease stages are displayed. Only species with interactions are shown. Co-exclusion interactions are displayed in red.

We can further investigate species differences by inferring prevalence differences between disease states of samples, using $\chi^2$ testing, tables S4-S8.

**Table S4:** prevalence differences between sample locations, SRP172818.

| species | association | pvalue | | normal | peripherical | tumor | count |
|---|---|---|---|---|---|---|---|
| Fusobacterium mortiferum | normal | 6.4e-05 | *** | 9/57 (15.8%) | 0/57 (0.0%) | 0/59 (0.0%) | 9 |
| Streptomyces atroolivaceus | normal | 3.2e-03 | ** | 7/57 (12.3%) | 1/57 (1.8%) | 0/59 (0.0%) | 8 |
| Peptostreptococcus stomatis | peripherical,tumor | 9.2e-03 | ** | 15/57 (26.3%) | 24/57 (42.1%) | 32/59 (54.2%) | 71 |
| Corynebacterium tuberculostearicum | tumor | 1.6e-04 | *** | 2/57 (3.5%) | 1/57 (1.8%) | 13/59 (22.0%) | 16 |
| Propionibacterium acnes | tumor | 3.5e-04 | *** | 25/57 (43.9%) | 22/57 (38.6%) | 43/59 (72.9%) | 90 |
| Bifidobacterium dentium | tumor | 5.6e-04 | *** | 2/57 (3.5%) | 3/57 (5.3%) | 14/59 (23.7%) | 19 |
| Rudaeicoccus suwonensis | tumor | 6.8e-04 | *** | 3/57 (5.3%) | 7/57 (12.3%) | 18/59 (30.5%) | 28 |
| Deinococcus citri | tumor | 1.6e-03 | ** | 1/57 (1.8%) | 0/57 (0.0%) | 8/59 (13.6%) | 9 |
| Campylobacter rectus | tumor | 1.6e-03 | ** | 1/57 (1.8%) | 0/57 (0.0%) | 8/59 (13.6%) | 9 |
| Actinomyces odontolyticus | tumor | 1.8e-03 | ** | 1/57 (1.8%) | 7/57 (12.3%) | 14/59 (23.7%) | 22 |
| Rothia mucilaginosa | tumor | 3.2e-03 | ** | 10/57 (17.5%) | 11/57 (19.3%) | 25/59 (42.4%) | 46 |
| Fusobacterium nucleatum | tumor | 5.7e-03 | ** | 8/57 (14.0%) | 12/57 (21.1%) | 23/59 (39.0%) | 43 |
| Fusobacterium periodonticum | tumor | 5.9e-03 | ** | 1/57 (1.8%) | 6/57 (10.5%) | 12/59 (20.3%) | 19 |
| Alloprevotella tannerae | tumor | 7.7e-03 | ** | 1/57 (1.8%) | 2/57 (3.5%) | 9/59 (15.3%) | 12 |
| Prevotella melaninogenica | tumor | 3.4e-02 | * | 2/57 (3.5%) | 4/57 (7.0%) | 10/59 (16.9%) | 16 |
| Helicobacter pylori | tumor | 4.4e-02 | * | 47/57 (82.5%) | 51/57 (89.5%) | 57/59 (96.6%) | 155 |
| Parvimonas micra | tumor | 7.3e-02 | | 15/57 (26.3%) | 14/57 (24.6%) | 25/59 (42.4%) | 54 |

**Table S5:** prevalence differences between sample locations, SRP128749.

| species | association | pvalue | | normal | peripherical | tumor | count |
|---|---|---|---|---|---|---|---|
| Fusobacterium mortiferum | normal | 0.0e+00 | *** | 25/225 (11.1%) | 3/215 (1.4%) | 1/229 (0.4%) | 29 |
| Bifidobacterium longum | normal | 0.0e+00 | *** | 42/225 (18.7%) | 4/215 (1.9%) | 18/229 (7.9%) | 64 |
| Clostridium cellulovorans | normal | 1.3e-05 | *** | 16/225 (7.1%) | 2/215 (0.9%) | 1/229 (0.4%) | 19 |
| Prevotella stercorea | normal | 7.1e-05 | *** | 22/225 (9.8%) | 2/215 (0.9%) | 9/229 (3.9%) | 33 |
| Nocardioides szechwanensis | normal | 6.7e-04 | *** | 9/225 (4.0%) | 1/215 (0.5%) | 0/229 (0.0%) | 10 |
| Roseburia inulinivorans | normal | 9.1e-04 | *** | 10/225 (4.4%) | 0/215 (0.0%) | 2/229 (0.9%) | 12 |
| Barnesiella intestinihominis | normal | 9.3e-04 | *** | 7/225 (3.1%) | 0/215 (0.0%) | 0/229 (0.0%) | 7 |
| Bacteroides uniformis | normal | 2.4e-03 | ** | 17/225 (7.6%) | 2/215 (0.9%) | 9/229 (3.9%) | 28 |
| Deinococcus aetherius | normal | 2.5e-03 | ** | 6/225 (2.7%) | 0/215 (0.0%) | 0/229 (0.0%) | 6 |
| Sulfurospirillum deleyianum | normal | 2.5e-03 | ** | 6/225 (2.7%) | 0/215 (0.0%) | 0/229 (0.0%) | 6 |
| Nitrospira japonica | normal | 4.4e-03 | ** | 11/225 (4.9%) | 2/215 (0.9%) | 2/229 (0.9%) | 15 |
| Parabacteroides merdae | normal | 6.9e-03 | ** | 5/225 (2.2%) | 0/215 (0.0%) | 0/229 (0.0%) | 5 |
| Ruminococcus bromii | normal,tumor | 7.0e-07 | *** | 57/225 (25.3%) | 14/215 (6.5%) | 42/229 (18.3%) | 113 |
| Faecalibacterium prausnitzii | normal,tumor | 3.7e-06 | *** | 127/225 (56.4%) | 71/215 (33.0%) | 111/229 (48.5%) | 309 |
| Arthrobacter oxydans | normal,tumor | 1.2e-03 | ** | 47/225 (20.9%) | 28/215 (13.0%) | 62/229 (27.1%) | 137 |
| Pyramidobacter piscolens | normal,tumor | 3.4e-03 | ** | 17/225 (7.6%) | 2/215 (0.9%) | 11/229 (4.8%) | 30 |
| Roseomonas gilardii | peripherical | 2.2e-03 | ** | 4/225 (1.8%) | 20/215 (9.3%) | 12/229 (5.2%) | 36 |
| Sphingomonas yabuuchiae | peripherical | 7.2e-03 | ** | 66/225 (29.3%) | 94/215 (43.7%) | 82/229 (35.8%) | 242 |
| Roseomonas stagni | peripherical | 9.1e-03 | ** | 4/225 (1.8%) | 17/215 (7.9%) | 10/229 (4.4%) | 31 |
| Helicobacter pylori | peripherical,tumor | 4.8e-02 | * | 155/225 (68.9%) | 169/215 (78.6%) | 175/229 (76.4%) | 499 |
| Peptostreptococcus stomatis | tumor | 0.0e+00 | *** | 52/225 (23.1%) | 60/215 (27.9%) | 130/229 (56.8%) | 242 |
| Propionibacterium acnes | tumor | 0.0e+00 | *** | 82/225 (36.4%) | 65/215 (30.2%) | 140/229 (61.1%) | 287 |
| Parvimonas micra | tumor | 3.0e-07 | *** | 39/225 (17.3%) | 40/215 (18.6%) | 85/229 (37.1%) | 164 |
| Fusobacterium nucleatum | tumor | 7.0e-07 | *** | 34/225 (15.1%) | 45/215 (20.9%) | 82/229 (35.8%) | 161 |
| Campylobacter showae | tumor | 6.4e-06 | *** | 0/225 (0.0%) | 1/215 (0.5%) | 14/229 (6.1%) | 15 |
| Sphingomonas faeni | tumor | 1.9e-05 | *** | 29/225 (12.9%) | 47/215 (21.9%) | 71/229 (31.0%) | 147 |
| Catonella morbi | tumor | 2.7e-05 | *** | 7/225 (3.1%) | 8/215 (3.7%) | 29/229 (12.7%) | 44 |
| Thermus scotoductus | tumor | 9.7e-05 | *** | 63/225 (28.0%) | 48/215 (22.3%) | 93/229 (40.6%) | 204 |
| Corynebacterium tuberculostearicum | tumor | 9.9e-05 | *** | 10/225 (4.4%) | 8/215 (3.7%) | 30/229 (13.1%) | 48 |
| Leptotrichia wadei | tumor | 2.3e-04 | *** | 13/225 (5.8%) | 20/215 (9.3%) | 40/229 (17.5%) | 73 |
| Gardnerella vaginalis | tumor | 7.6e-04 | *** | 2/225 (0.9%) | 1/215 (0.5%) | 12/229 (5.2%) | 15 |
| Corynebacterium mucifaciens | tumor | 1.7e-03 | ** | 4/225 (1.8%) | 10/215 (4.7%) | 21/229 (9.2%) | 35 |
| Filifactor alocis | tumor | 4.3e-03 | ** | 9/225 (4.0%) | 13/215 (6.0%) | 27/229 (11.8%) | 49 |

**Table S6:** prevalence differences between disease stages, SRP070925.

| species | association | pvalue | | advanced.cancer | early.cancer | gastritis | metaplasia | count |
|---|---|---|---|---|---|---|---|---|
| Peptostreptococcus stomatis | advanced cancer | 2.6e-01 | | 3/20 (15.0%) | 0/20 (0.0%) | 1/20 (5.0%) | 1/20 (5.0%) | 5 |
| Novosphingobium sediminicola | early cancer,advanced cancer | 0.0e+00 | *** | 15/20 (75.0%) | 17/20 (85.0%) | 0/20 (0.0%) | 6/20 (30.0%) | 38 |
| Methylobacterium populi | gastritis | 1.5e-03 | ** | 7/20 (35.0%) | 6/20 (30.0%) | 17/20 (85.0%) | 8/20 (40.0%) | 38 |
| Sphingobium amiense | gastritis | 5.5e-03 | ** | 0/20 (0.0%) | 0/20 (0.0%) | 4/20 (20.0%) | 0/20 (0.0%) | 4 |
| Burkholderia cepacia | gastritis | 6.7e-03 | ** | 1/20 (5.0%) | 0/20 (0.0%) | 6/20 (30.0%) | 1/20 (5.0%) | 8 |
| Sphingomonas hunanensis | gastritis | 7.3e-03 | ** | 1/20 (5.0%) | 1/20 (5.0%) | 8/20 (40.0%) | 3/20 (15.0%) | 13 |
| Parvimonas micra | gastritis,advanced cancer | 3.1e-01 | | 3/20 (15.0%) | 1/20 (5.0%) | 2/20 (10.0%) | 0/20 (0.0%) | 6 |
| Sphingomonas faeni | gastritis,early cancer | 8.1e-03 | ** | 2/20 (10.0%) | 5/20 (25.0%) | 10/20 (50.0%) | 2/20 (10.0%) | 19 |
| Hyphomonas polymorpha | gastritis,metaplasia | 8.0e-07 | *** | 4/20 (20.0%) | 0/20 (0.0%) | 16/20 (80.0%) | 7/20 (35.0%) | 27 |
| Modestobacter multiseptatus | gastritis,metaplasia | 1.9e-04 | *** | 0/20 (0.0%) | 1/20 (5.0%) | 10/20 (50.0%) | 7/20 (35.0%) | 18 |
| Paenibacillus humicus | gastritis,metaplasia | 5.1e-03 | ** | 10/20 (50.0%) | 12/20 (60.0%) | 18/20 (90.0%) | 18/20 (90.0%) | 58 |
| Geotoga petraea | gastritis,metaplasia | 8.3e-03 | ** | 3/20 (15.0%) | 0/20 (0.0%) | 7/20 (35.0%) | 8/20 (40.0%) | 18 |
| Prevotella melaninogenica | gastritis,metaplasia | 4.5e-01 | | 3/20 (15.0%) | 4/20 (20.0%) | 6/20 (30.0%) | 7/20 (35.0%) | 20 |
| Helicobacter pylori | gastritis,metaplasia | 5.6e-01 | | 13/20 (65.0%) | 12/20 (60.0%) | 14/20 (70.0%) | 16/20 (80.0%) | 55 |
| Fusobacterium nucleatum | metaplasia,advanced cancer | 1.6e-01 | | 8/20 (40.0%) | 4/20 (20.0%) | 2/20 (10.0%) | 5/20 (25.0%) | 19 |

**Table S7:** prevalence differences between disease stages, ERP023334.

| species | association | pvalue | | cancer | dysplasia | gastritis | healthy | metaplasia | count |
|---|---|---|---|---|---|---|---|---|---|
| Lactobacillus rhamnosus | cancer | 2.0e-03 | ** | 2/10 (20.0%) | 0/8 (0.0%) | 0/44 (0.0%) | 0/22 (0.0%) | 0/9 (0.0%) | 2 |
| Corynebacterium pseudodiphtheriticum | dysplasia | 5.0e-07 | *** | 0/10 (0.0%) | 4/8 (50.0%) | 0/44 (0.0%) | 1/22 (4.5%) | 0/9 (0.0%) | 5 |
| Marmoricola aequoreus | dysplasia | 2.3e-04 | *** | 0/10 (0.0%) | 2/8 (25.0%) | 0/44 (0.0%) | 0/22 (0.0%) | 0/9 (0.0%) | 2 |
| Parvimonas micra | gastritis | 6.9e-01 | | 0/10 (0.0%) | 0/8 (0.0%) | 2/44 (4.5%) | 0/22 (0.0%) | 0/9 (0.0%) | 2 |
| Prevotella fusca | healthy | 1.1e-05 | *** | 0/10 (0.0%) | 0/8 (0.0%) | 0/44 (0.0%) | 8/22 (36.4%) | 0/9 (0.0%) | 8 |
| Haemophilus sputorum | healthy | 3.6e-04 | *** | 0/10 (0.0%) | 0/8 (0.0%) | 0/44 (0.0%) | 6/22 (27.3%) | 0/9 (0.0%) | 6 |
| Prevotella veroralis | healthy | 4.8e-04 | *** | 1/10 (10.0%) | 1/8 (12.5%) | 6/44 (13.6%) | 13/22 (59.1%) | 1/9 (11.1%) | 22 |
| Prevotella loescheii | healthy | 6.6e-04 | *** | 3/10 (30.0%) | 1/8 (12.5%) | 9/44 (20.5%) | 15/22 (68.2%) | 1/9 (11.1%) | 29 |
| Prevotella dentalis | healthy | 1.2e-03 | ** | 1/10 (10.0%) | 1/8 (12.5%) | 2/44 (4.5%) | 9/22 (40.9%) | 0/9 (0.0%) | 13 |
| Treponema amylovorum | healthy | 1.6e-03 | ** | 0/10 (0.0%) | 0/8 (0.0%) | 3/44 (6.8%) | 8/22 (36.4%) | 0/9 (0.0%) | 11 |
| Prevotella oulorum | healthy | 1.6e-03 | ** | 3/10 (30.0%) | 3/8 (37.5%) | 12/44 (27.3%) | 17/22 (77.3%) | 2/9 (22.2%) | 37 |
| Tannerella forsythia | healthy | 3.8e-03 | ** | 3/10 (30.0%) | 3/8 (37.5%) | 13/44 (29.5%) | 16/22 (72.7%) | 1/9 (11.1%) | 36 |
| Prevotella pallens | healthy | 4.2e-03 | ** | 3/10 (30.0%) | 3/8 (37.5%) | 17/44 (38.6%) | 17/22 (77.3%) | 1/9 (11.1%) | 41 |
| Treponema denticola | healthy | 4.6e-03 | ** | 2/10 (20.0%) | 1/8 (12.5%) | 6/44 (13.6%) | 11/22 (50.0%) | 0/9 (0.0%) | 20 |
| Propionibacterium acnes | healthy | 5.1e-02 | | 1/10 (10.0%) | 2/8 (25.0%) | 8/44 (18.2%) | 11/22 (50.0%) | 2/9 (22.2%) | 24 |
| Porphyromonas endodontalis | healthy,cancer | 1.5e-05 | *** | 4/10 (40.0%) | 2/8 (25.0%) | 9/44 (20.5%) | 19/22 (86.4%) | 3/9 (33.3%) | 37 |
| Alloprevotella rava | healthy,cancer | 3.7e-05 | *** | 4/10 (40.0%) | 2/8 (25.0%) | 8/44 (18.2%) | 17/22 (77.3%) | 1/9 (11.1%) | 32 |
| Solobacterium moorei | healthy,cancer | 6.7e-04 | *** | 3/10 (30.0%) | 2/8 (25.0%) | 6/44 (13.6%) | 13/22 (59.1%) | 0/9 (0.0%) | 24 |
| Neisseria elongata | healthy,dysplasia | 2.1e-05 | *** | 1/10 (10.0%) | 3/8 (37.5%) | 9/44 (20.5%) | 17/22 (77.3%) | 1/9 (11.1%) | 31 |
| Haemophilus parainfluenzae | healthy,dysplasia | 5.8e-05 | *** | 4/10 (40.0%) | 8/8 (100.0%) | 17/44 (38.6%) | 19/22 (86.4%) | 2/9 (22.2%) | 50 |
| Prevotella oris | healthy,dysplasia | 1.2e-04 | *** | 2/10 (20.0%) | 5/8 (62.5%) | 12/44 (27.3%) | 18/22 (81.8%) | 2/9 (22.2%) | 39 |
| Selenomonas dianae | healthy,dysplasia | 1.5e-04 | *** | 3/10 (30.0%) | 4/8 (50.0%) | 12/44 (27.3%) | 18/22 (81.8%) | 1/9 (11.1%) | 38 |
| Lautropia mirabilis | healthy,dysplasia | 2.0e-04 | *** | 2/10 (20.0%) | 5/8 (62.5%) | 10/44 (22.7%) | 16/22 (72.7%) | 1/9 (11.1%) | 34 |
| Actinomyces odontolyticus | healthy,dysplasia | 6.9e-04 | *** | 5/10 (50.0%) | 8/8 (100.0%) | 18/44 (40.9%) | 19/22 (86.4%) | 4/9 (44.4%) | 54 |
| Bradyrhizobium elkanii | healthy,dysplasia | 7.8e-04 | *** | 0/10 (0.0%) | 3/8 (37.5%) | 0/44 (0.0%) | 3/22 (13.6%) | 0/9 (0.0%) | 6 |
| Capnocytophaga gingivalis | healthy,dysplasia | 8.3e-04 | *** | 2/10 (20.0%) | 4/8 (50.0%) | 8/44 (18.2%) | 15/22 (68.2%) | 2/9 (22.2%) | 31 |
| Prevotella intermedia | healthy,dysplasia | 1.0e-03 | ** | 1/10 (10.0%) | 2/8 (25.0%) | 6/44 (13.6%) | 12/22 (54.5%) | 0/9 (0.0%) | 21 |
| Alloprevotella tannerae | healthy,dysplasia | 1.6e-03 | ** | 2/10 (20.0%) | 4/8 (50.0%) | 16/44 (36.4%) | 17/22 (77.3%) | 1/9 (11.1%) | 40 |
| Campylobacter curvus | healthy,dysplasia | 1.9e-03 | ** | 4/10 (40.0%) | 6/8 (75.0%) | 17/44 (38.6%) | 19/22 (86.4%) | 3/9 (33.3%) | 49 |
| Actinomyces graevenitzii | healthy,dysplasia | 1.9e-03 | ** | 1/10 (10.0%) | 5/8 (62.5%) | 12/44 (27.3%) | 15/22 (68.2%) | 2/9 (22.2%) | 35 |
| Prevotella salivae | healthy,dysplasia | 1.9e-03 | ** | 5/10 (50.0%) | 7/8 (87.5%) | 21/44 (47.7%) | 19/22 (86.4%) | 2/9 (22.2%) | 54 |
| Aggregatibacter segnis | healthy,dysplasia | 2.0e-03 | ** | 0/10 (0.0%) | 2/8 (25.0%) | 7/44 (15.9%) | 12/22 (54.5%) | 1/9 (11.1%) | 22 |
| Veillonella atypica | healthy,dysplasia | 2.9e-03 | ** | 4/10 (40.0%) | 7/8 (87.5%) | 16/44 (36.4%) | 15/22 (68.2%) | 1/9 (11.1%) | 43 |
| Porphyromonas catoniae | healthy,dysplasia | 3.1e-03 | ** | 2/10 (20.0%) | 5/8 (62.5%) | 16/44 (36.4%) | 17/22 (77.3%) | 2/9 (22.2%) | 42 |
| Streptococcus salivarius | healthy,dysplasia | 3.7e-03 | ** | 3/10 (30.0%) | 7/8 (87.5%) | 20/44 (45.5%) | 17/22 (77.3%) | 2/9 (22.2%) | 49 |
| Veillonella parvula | healthy,dysplasia | 5.0e-03 | ** | 3/10 (30.0%) | 6/8 (75.0%) | 15/44 (34.1%) | 15/22 (68.2%) | 1/9 (11.1%) | 40 |
| Streptococcus parasanguinis | healthy,dysplasia,cancer | 4.1e-05 | *** | 7/10 (70.0%) | 8/8 (100.0%) | 16/44 (36.4%) | 20/22 (90.9%) | 4/9 (44.4%) | 55 |
| Neisseria bacilliformis | healthy,dysplasia,cancer | 1.5e-04 | *** | 2/10 (20.0%) | 2/8 (25.0%) | 2/44 (4.5%) | 11/22 (50.0%) | 0/9 (0.0%) | 17 |
| Atopobium parvulum | healthy,dysplasia,cancer | 4.5e-03 | ** | 3/10 (30.0%) | 3/8 (37.5%) | 7/44 (15.9%) | 12/22 (54.5%) | 0/9 (0.0%) | 25 |
| Fusobacterium nucleatum | healthy,dysplasia,cancer | 7.5e-03 | ** | 8/10 (80.0%) | 7/8 (87.5%) | 25/44 (56.8%) | 18/22 (81.8%) | 2/9 (22.2%) | 60 |
| Sphingomonas paucimobilis | metaplasia | 7.6e-04 | *** | 0/10 (0.0%) | 0/8 (0.0%) | 0/44 (0.0%) | 0/22 (0.0%) | 2/9 (22.2%) | 2 |

Revised manuscript

**Table S8:** prevalence differences between disease stages, ERP023334.

| species | association | pvalue | | functional.dyspepsia | gastric.cancer | gastric.ulcer | count |
|---|---|---|---|---|---|---|---|
| Lachnoanaerobaculum umeaense | functional dyspepsia | 7.0e-03 | ** | 2/6 (33.3%) | 0/15 (0.0%) | 0/13 (0.0%) | 2 |
| Aquabacterium parvum | functional dyspepsia | 7.0e-03 | ** | 2/6 (33.3%) | 0/15 (0.0%) | 0/13 (0.0%) | 2 |
| Methylobacterium radiotolerans | functional dyspepsia,gastric ulcer | 1.3e-04 | *** | 6/6 (100.0%) | 5/15 (33.3%) | 13/13 (100.0%) | 24 |
| Lactococcus lactis | gastric cancer | 5.2e-04 | *** | 2/6 (33.3%) | 12/15 (80.0%) | 1/13 (7.7%) | 15 |
| Comamonas testosteroni | gastric cancer | 9.9e-03 | ** | 0/6 (0.0%) | 6/15 (40.0%) | 0/13 (0.0%) | 6 |
| Peptostreptococcus stomatis | gastric cancer | 2.6e-01 | | 0/6 (0.0%) | 2/15 (13.3%) | 0/13 (0.0%) | 2 |
| Parvimonas micra | gastric cancer | 5.2e-01 | | 0/6 (0.0%) | 1/15 (6.7%) | 0/13 (0.0%) | 1 |
| Fusobacterium nucleatum | gastric cancer,gastric ulcer | 6.4e-01 | | 1/6 (16.7%) | 5/15 (33.3%) | 5/13 (38.5%) | 11 |

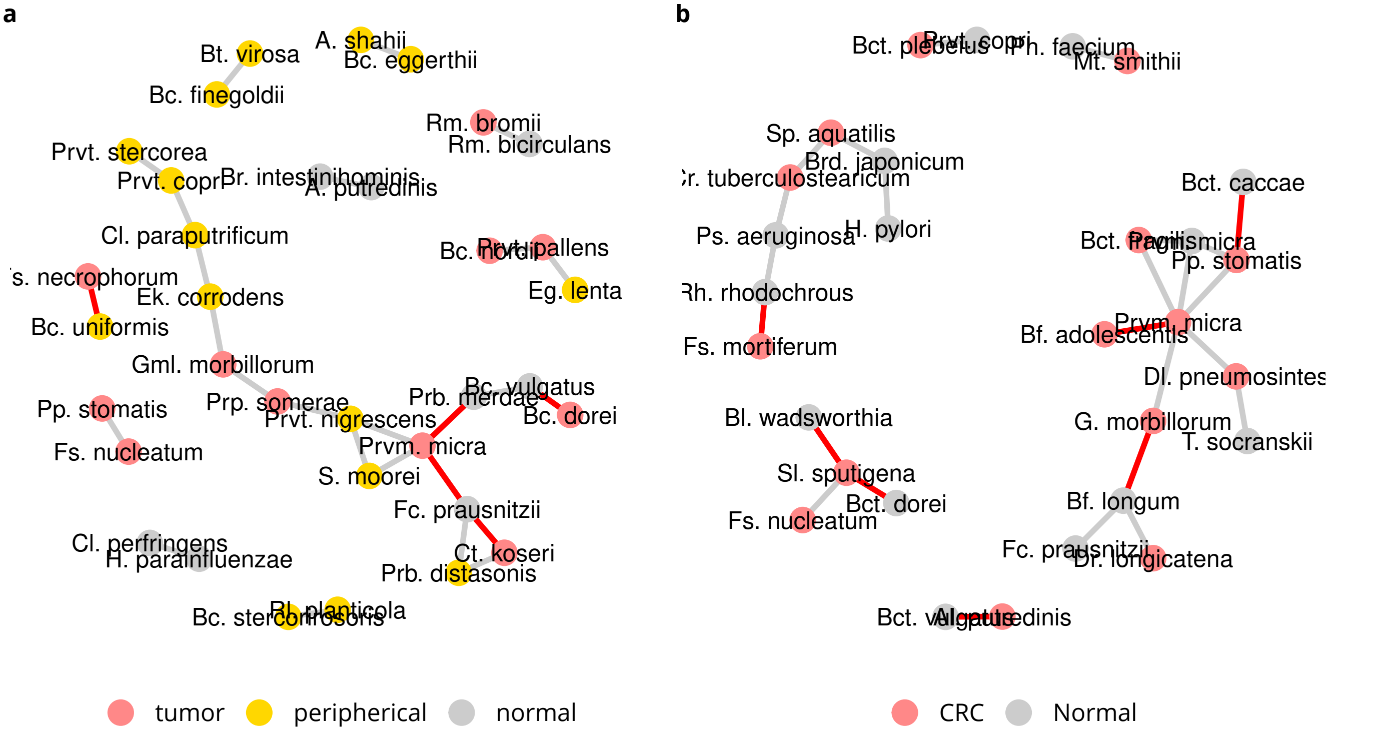

**Figure S12:** Discriminating species in CRC. Data sets a) SRP137015 and b) SRP076561. Only species with interactions are displayed. Location associations are based on maximum mean relative abundance. Co-exclusion is indicated in red.

#### comparison with CRC

We test two CRC data sets for presence and interactions of *F. nucleatum*, *P. micra* and *P. stomatis*. Data set SRP117763 (n=34, tumor-only) was published by [Purcell et al. 2017] and data set SRP137015 (n=211, tumor/peripherical/normal) by [Hale et al. 2018b;a]. We find *F. nucleatum* in interaction with *P. stomatis* in SRP137015 and *P. micra* in interaction with *P. stomatis* in SRP117763, figure S12. Prevalence of *F. nucleatum* is over 70% in tumor samples in SRP117763, table S9 and at 48% in SRP137015, table S10.

**Table S9:** prevalence differences between CRC subtypes, SRP117763.

| species | association | pvalue | | CMS1 | CMS2 | CMS3 | count |
|---|---|---|---|---|---|---|---|
| Clostridium cadaveris | CMS1 | 5.1e-03 | ** | 4/6 (66.7%) | 2/13 (15.4%) | 0/10 (0.0%) | 6 |
| Parvimonas micra | CMS1,CMS2 | 4.0e-02 | * | 3/6 (50.0%) | 8/13 (61.5%) | 1/10 (10.0%) | 12 |
| Peptostreptococcus stomatis | CMS1,CMS2 | 1.8e-01 | | 2/6 (33.3%) | 6/13 (46.2%) | 1/10 (10.0%) | 9 |
| Prevotella melaninogenica | CMS1,CMS2 | 4.4e-01 | | 1/6 (16.7%) | 1/13 (7.7%) | 0/10 (0.0%) | 2 |
| Fusobacterium nucleatum | CMS1,CMS2 | 8.3e-01 | | 5/6 (83.3%) | 10/13 (76.9%) | 7/10 (70.0%) | 22 |

**Table S10:** prevalence differences between CRC sample locations, SRP137015.

| species | association | pvalue | | normal | peripherical | tumor | count |
|---|---|---|---|---|---|---|---|
| Prevotella melaninogenica | normal | 5.9e-01 | | 1/103 (1.0%) | 0/46 (0.0%) | 0/62 (0.0%) | 1 |
| Bacteroides vulgatus | normal,peripherical | 1.3e-03 | ** | 80/103 (77.7%) | 38/46 (82.6%) | 34/62 (54.8%) | 152 |
| Peptostreptococcus stomatis | peripherical,tumor | 1.7e-01 | | 12/103 (11.7%) | 8/46 (17.4%) | 14/62 (22.6%) | 34 |
| Fusobacterium nucleatum | tumor | 3.4e-04 | *** | 20/103 (19.4%) | 12/46 (26.1%) | 30/62 (48.4%) | 62 |
| Campylobacter gracilis | tumor | 1.5e-03 | ** | 1/103 (1.0%) | 1/46 (2.2%) | 8/62 (12.9%) | 10 |
| Parvimonas micra | tumor | 2.4e-03 | ** | 5/103 (4.9%) | 5/46 (10.9%) | 14/62 (22.6%) | 24 |

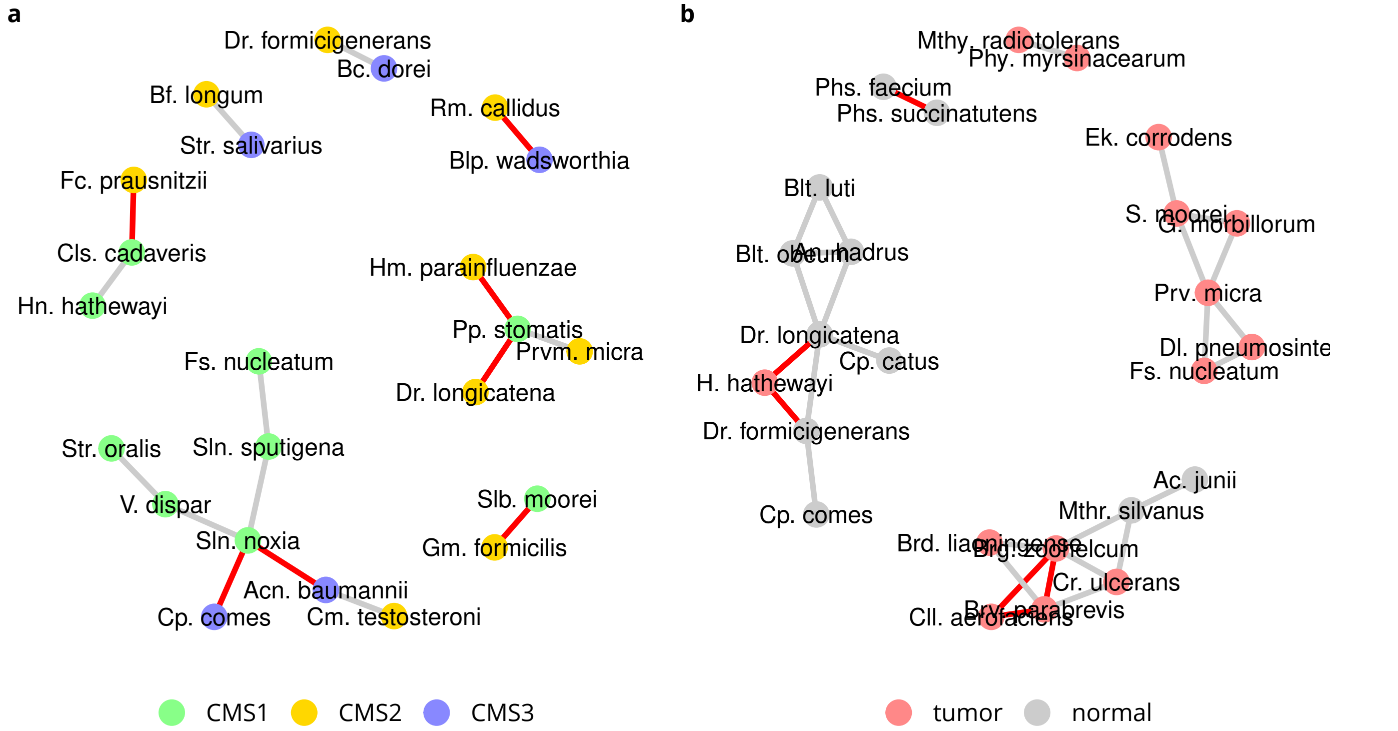

**Figure S13:** Discriminating species in CRC. Data sets a) SRP117763 and b) ERP005534 Only species with interactions are displayed. Location associations are based on maximum mean relative abundance. Co-exclusion is indicated in red.

**Table S11:** prevalence differences between CRC sample locations, SRP076561.

| species | association | pvalue | CRC | Normal | count |
|---|---|---|---|---|---|
| Fusobacterium nucleatum | tumor | 0.17 | 19/26 (73.1%) | 12/24 (50.0%) | 31 |
| Prevotella melaninogenica | tumor | 1.00 | 1/26 (3.8%) | 0/24 (0.0%) | 1 |
| Propionibacterium acnes | normal | 0.17 | 8/26 (30.8%) | 13/24 (54.2%) | 21 |
| Parvimonas micra | normal | 0.72 | 15/26 (57.7%) | 16/24 (66.7%) | 31 |
| Helicobacter pylori | normal | 0.74 | 14/26 (53.8%) | 15/24 (62.5%) | 29 |
| Peptostreptococcus stomatis | normal | 1.00 | 16/26 (61.5%) | 15/24 (62.5%) | 31 |

**Table S12:** prevalence differences between CRC sample locations, ERP005534.

| species | association | pvalue | normal | tumor | count |
|---|---|---|---|---|---|
| Parvimonas micra | | 1.00 | 33/48 (68.8%) | 33/48 (68.8%) | 66 |
| Prevotella melaninogenica | normal | 0.47 | 2/48 (4.2%) | 0/48 (0.0%) | 2 |
| Fusobacterium nucleatum | tumor | 0.11 | 31/48 (64.6%) | 39/48 (81.2%) | 70 |
| Peptostreptococcus stomatis | tumor | 0.68 | 22/48 (45.8%) | 25/48 (52.1%) | 47 |

#### H. pylori proportion

In dataset SRP070925 high (>50%) *H. pylori* proportion is not detected, contrary to the healthy cohort SRP200169. In datasets SRP128749 and SRP172818, high *H. pylori* proportions are detected in subsets of all three micro-environments normal, peripherical and tumor. *H. pylori* free samples are found in all datasets, indistinctive of disease progress or micro-environment.

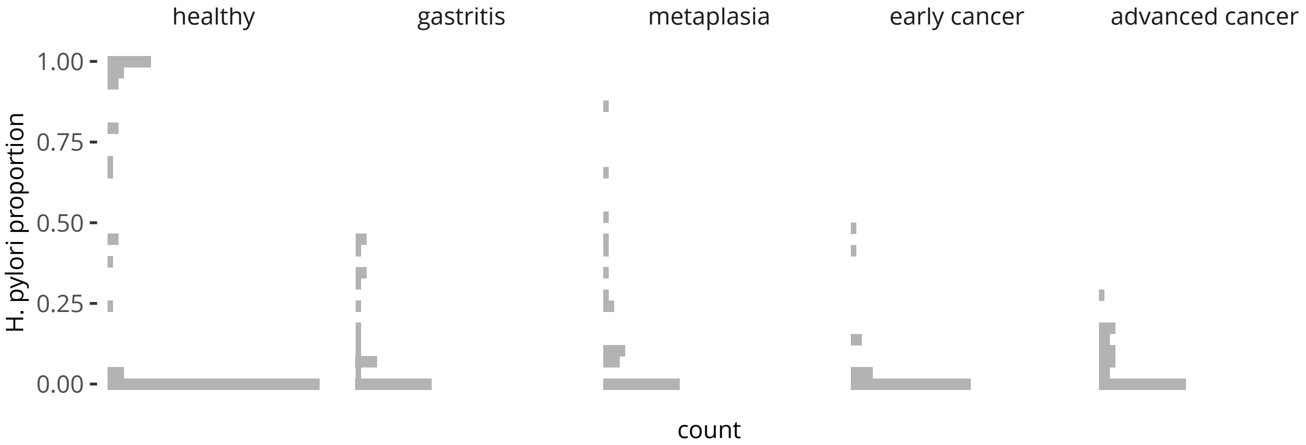

**Figure S14:** Helicobacter pylori proportion, SRP200169 + SRP070925.

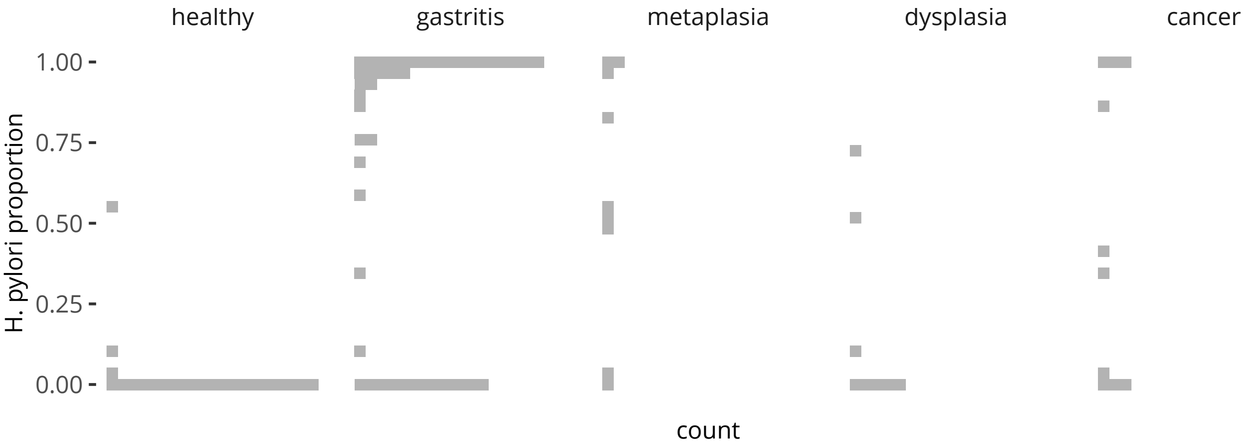

**Figure S15:** Helicobacter pylori proportion, ERP023334.

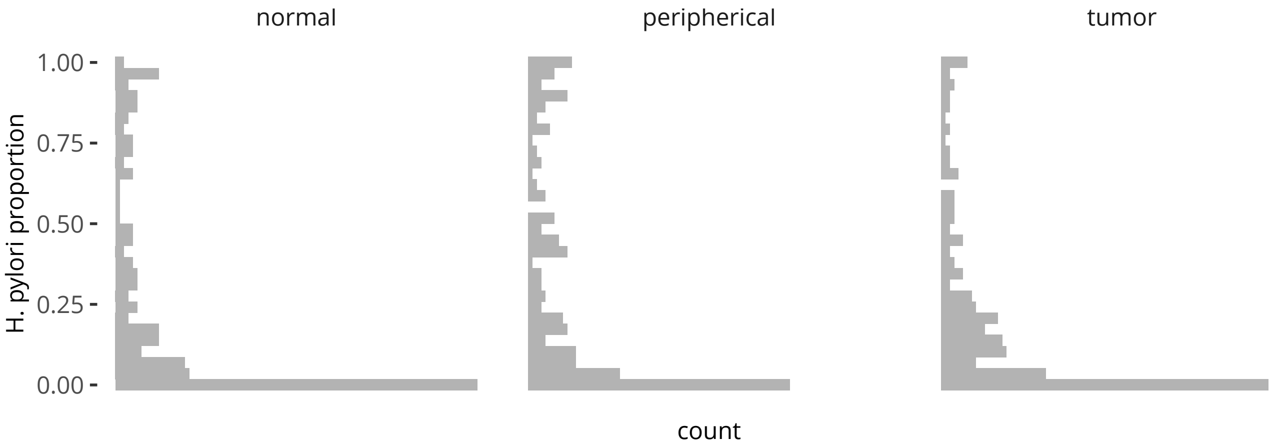

**Figure S16:** Helicobacter pylori proportion, SRP128749.

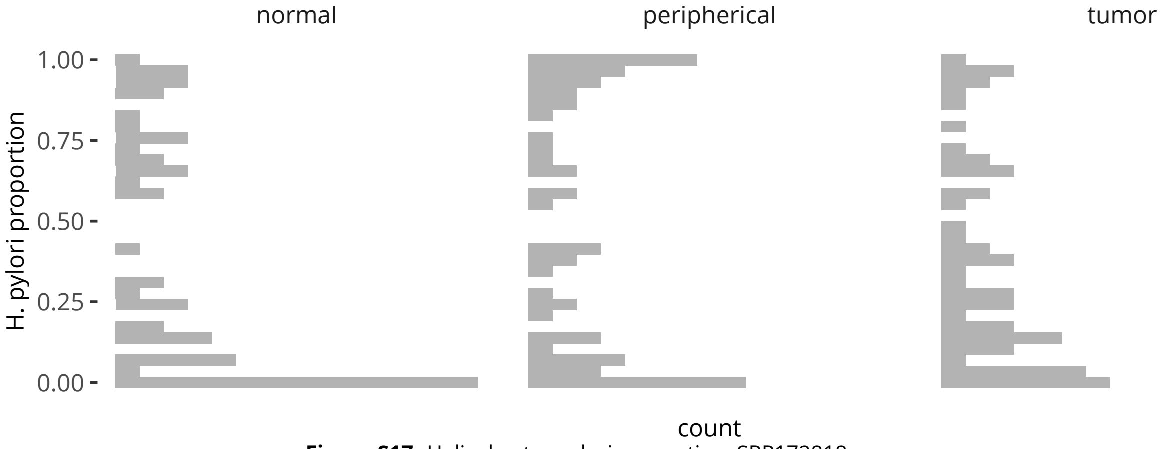

**Figure S17:** Helicobacter pylori proportion, SRP172818.
